# Nitrocellulose redox permanganometry: a simple method for reductive capacity assessment

**DOI:** 10.1101/2020.06.16.154682

**Authors:** J Homolak, I Kodvanj, A Babic Perhoc, D Virag, A Knezovic, J Osmanovic Barilar, P Riederer, M Salkovic-Petrisic

## Abstract

We propose a rapid, simple and robust method for measurement of reductive capacity of liquid and solid biological samples based on potassium permanganate reduction followed by trapping of manganese dioxide precipitate on a nitrocellulose membrane. Moreover, we discuss how nitrocellulose redox permanganometry (NRP) can be used for high-throughput analysis of biological samples and present HistoNRP, its modification used for detailed analysis of reductive capacity spatial distribution in tissue with preserved anatomical relations.

## Main text

Under physiological conditions, cellular homeostatic machinery keeps the production and elimination of biological free radicals in balance. However, once this balance is impaired, unconstrained accumulation of free radicals triggers pathophysiological cascades involved in the etiopathogenesis of a myriad of disorders^1–3^. Consequently, numerous methods have been developed in order to study the production and elimination of free radicals in biological samples. The complexity of the redox homeostatic system and problematic analysis of individual subsystems lead to the need to develop a concept of a single test that might reflect total reductive capacity and thus provide information on the overall redox status of the sample. This information is invaluable for oxidative stress research as it often provides context for understanding whether the changes of systems involved in redox homeostasis are of etiopathogenetic or compensatory nature. Hence, total reductive capacity, often called total antioxidant capacity (TAC), has been one of the most widely used methods in oxidative stress research. Consequently, many assays have been developed in order to determine TAC of biological samples, but not without limitations^4^. Most of the TAC assays are based on complex chemical reactions, require sophisticated equipment and complex sample processing techniques that can potentially introduce experimental bias. Additionally, standard methods are often time-consuming, require a relatively large amount of biological samples and, most importantly, none of the existing methods can be used for analysis of both liquid (e.g. plasma, serum, tissue homogenate) and solid (e.g. formalin-fixed paraffin-embedded or frozen tissue samples) biological samples, let alone provide the information on anatomical distribution of reduction potentials.

Here, we propose a rapid, simple and robust method for measurement of reductive capacity suitable for high-throughput sample processing. The proposed method can be used to understand the anatomical distribution of tissue reductive capacity, something we believe is an invaluable piece of information in oxidative stress research. In our method, reductive capacity of the sample is measured by its capacity to reduce potassium permanganate (KMnO_4_) to manganese dioxide (MnO_2_) in a neutral environment, but in contrast to other methods based on potassium permanganate reduction, the reaction takes place on a nitrocellulose membrane. Potassium permanganate is commonly used for redox titrations ever since the discovery of permanganometry by Margueritte in 1846^5^. It is a strong oxidant, enters few non-redox reactions and is therefore characterized as an ideal reactant for TAC determinations^4^. However, its use in classic redox titrations is limited by an abundant brown precipitate of MnO_2_ generated by KMnO_4_ reduction in a neutral environment that complicates determination of the reaction endpoint^4^. In the proposed method, we used this limitation to our advantage by trapping the precipitated brown MnO_2_ during the redox reaction in a nitrocellulose membrane and using the solid precipitate for quantification of sample-mediated locally reduced KMnO_4_. MnO_2_ can be used for precise quantification of reductive capacity, as previously elegantly shown by Zhou et al. ^4^, where increased concentrations of MnO_2_ in potassium permanganate agar were used to determine the antioxidant capacity of human sera.

The schematic overview of our method is depicted in **Fig 1A** and a detailed step-by-step protocol is available in **Supplement 1**. In order to test linearity, accuracy and precision of the proposed method, named nitrocellulose redox permanganometry (NRP), we used sodium thiosulfate (Na_2_S_2_O_3_), a well-known reducing agent standardly used for TAC method validation. Briefly, we independently prepared 10 stock solutions with nominal concentrations of 0.1M – 0.01M (**Fig 1B**) (and explained in detail in the **Supplement 2**) and tested the method by using the validation criteria adopted from the official European Medicines Agency Bioanalytical Method Validation guidelines^6^. To assure the results were not specific for Na_2_S_2_O_3_ and that they truly reflected antioxidant capacity, the whole procedure was repeated with another standard reducing agent, ascorbic acid (AA) (**Fig 1C**). Graded nominal concentrations of classic antioxidants are standardly used for validation of TAC methods. However, we believe graded standards suffer from vulnerability to chemical concentration bias, so we introduced an additional test with physically oxidized isoconcentrated samples of AA (**Fig 1D**). Further validation of the observed effect was done by time and temperature response analysis of AA oxidation (**Supplement 3**). As results of both validation protocols suggested NRP to be a robust and precise method for reduction capacity assessment, additional tests were done to identify an optimal signal quantification protocol. We analyzed NRP membrane stability (**Supplement 4**) as well as potential differences in membrane digitalization techniques that could affect signal quantification (**Supplement 5**). Regarding the computational analysis of signal intensity, a method relying on the Fiji (NIH, USA) Gel Analyzer plugin was determined to be the best. The protocol is explained in **Supplement 6**. After all validation standards were fulfilled, we wanted to verify whether the same principles demonstrated with simple chemical samples can also be applied to chemically complex biological systems. To do this, we measured the direct oxidation-reduction potential (ORP) of graded nominal concentrations of Na2S2O3 by a redox microsensor system composed of a platinum sensing element and an Ag/AgCl reference, and compared these measurements to NRP results of the same Na_2_S_2_O_3_ samples (**Fig 1E**). We then used these samples for exogenous manipulation of one biological specimen (hippocampal homogenate) to create an experimental set with subtle reduction potential grading. We proceeded to test this experimental set by both ORP and NRP measurements (**Fig 1F**).

**Fig. 1.**
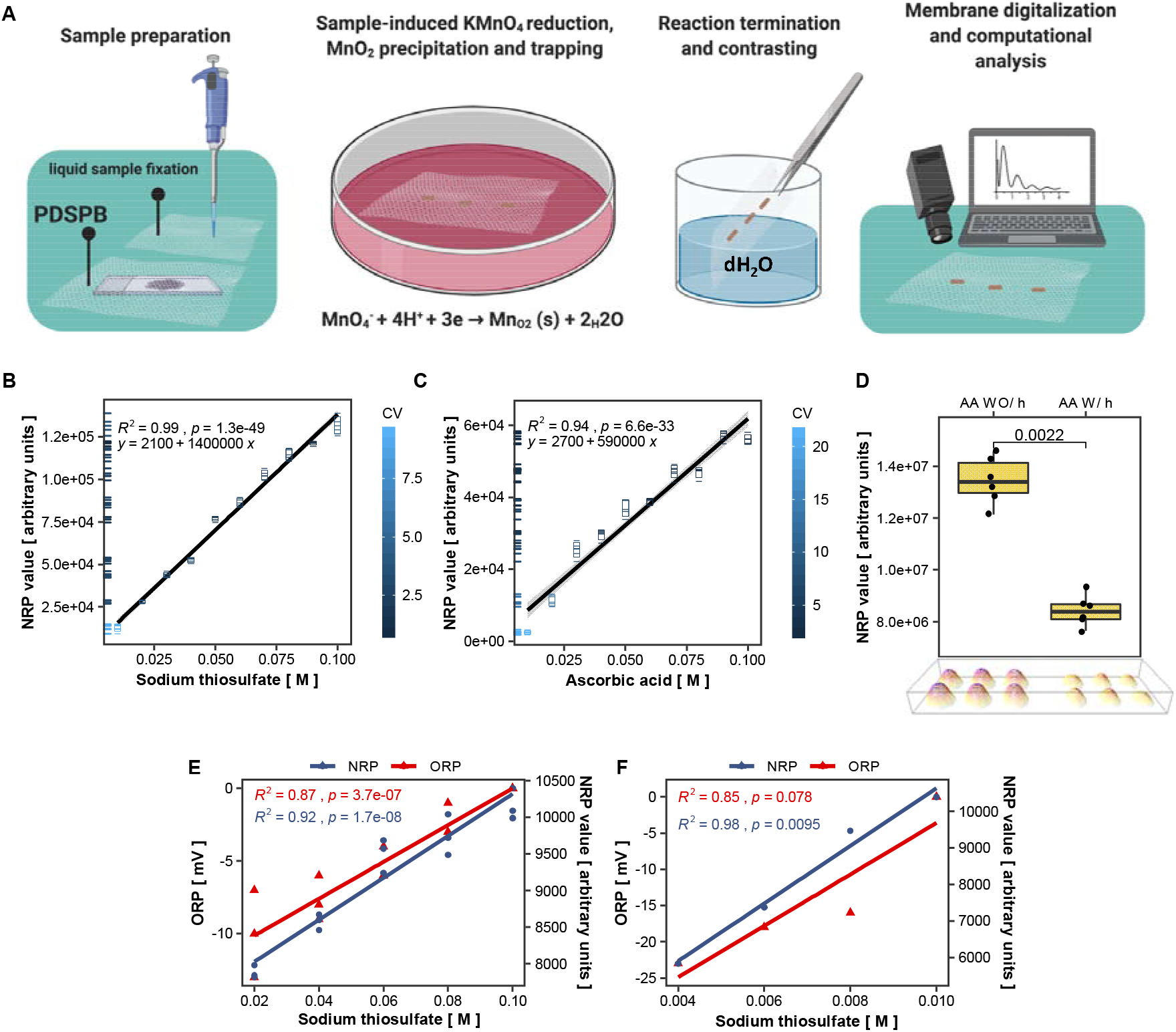
Validation of nitrocellulose redox permanganometry (NRP). **A**) Schematic step-by-step overview of NRP. First, samples of interest are fixed onto a clean sheet of nitrocellulose. Liquid samples (chemical solutions, serum, plasma, tissue homogenates) are fixed onto the membrane by pipetting 1 μl of sample and leaving the membrane to air-dry. Solid samples (formalin-fixed paraffin embedded (FFPE) tissue samples and cryosections) are fixed onto nitrocellulose by enzymatic retrieval followed by heat-facilitated passive diffusion slice printing for FFPE or passive diffusion slice print blotting (PDSPB) for cryosections (described in detail in the **Supplement 1**). After drying, the membrane is immersed in a potassium permanganate solution (63.27 mM in ddH_2_O) for 30 seconds. The reaction is terminated by placing the membrane in dH_2_O until the contrast is maximized and the membrane is left to dry out. Once dry, the nitrocellulose sheet is digitalized and analyzed. **B**) Validation of NRP by Na_2_S_2_O_3_. Ten stock solutions of Na_2_S_2_O_3_ (0.01-0.1 M) were independently prepared and fixed onto a nitrocellulose membrane. Each concentration was applied to the membrane 5-6 times and the membrane was analyzed by the protocol described in A). The integrated intensity of all dots was calculated in Fiji and statistically analyzed and visualized in R. Accuracy, precision and linearity were calculated and are available in the **Supplement 2**. All points were colored by coefficient of variation (CV) for respective concentrations. Individual data points were additionally represented on the y axis by ticks color-coded by their CV. **C**) Validation of NRP by ascorbic acid. In short, the whole procedure described in B) was repeated with independently prepared stock solutions of ascorbic acid (0.01-0.1 M). **D**) Validation of NRP with physically oxidized isoconcentrated samples of ascorbic acid. Briefly, we independently prepared six solutions of 0.05 M ascorbic acid and aliquoted each sample in two Eppendorf tubes. One of two aliquots was placed at 4°C and the other in a heating block at 70°C. After 4 hours, both aliquots were left to acclimate to room temperature and then 1 μl of each solution was placed on a clean sheet of nitrocellulose membrane. Once dry, the membrane was analyzed by the protocol described in A). Integrated intensity of all dots was calculated in Fiji and statistically analyzed and visualized in R. Fire lookup table 3D intensity surface plot of raw images is presented at the bottom of the graph. Ascorbic acid oxidation validation was additionally tested in respect to time and temperature (all experiments are available in the **Supplement 3**). AA – ascorbic acid; WO/h – without heating; W/h – with heating. **E**) Comparison of platinum microelectrode-measured direct electrochemical oxidation-reduction potential (ORP) with NRP. Five graded nominal concentrations of Na_2_S_2_O_3_ were prepared and measured with ORP and NRP. Both tests were performed in triplicates (additional replications are available in **Supplement 7**). Coefficient of determination (R^2^) and p values were calculated for both methods. Statistical analysis and visualization was conducted with R. **F**) The tested nominal concentrations of Na_2_S_2_O_3_ (from E) were used for exogenous redox manipulation of a single biological specimen (hippocampal homogenate). In short, 9 μl of tissue homogenate was aliquoted in separate tubes, vortexed, spun down and treated with 1 μl of ddH_2_O or one of the nominal solutions of Na_2_S_2_O_3_ described in the previous experiment. All samples were vortexed, spun down and analyzed with ORP and NRP to compare linearity and sensitivity of both methods to slight redox manipulations in a biological system. Coefficient of determination was calculated for both methods (additional replications are available in the **Supplement 7**).

After showing that NRP is compatible with reductive capacity measurements of biological specimens, we moved on to demonstrate the method on animal samples from our previous experiments (described in detail in the **Supplement 8**). Here, we used two common models of Alzheimer’s disease, a rat model of sporadic Alzheimer’s disease (sAD) induced by intracerebroventricular streptozotocin (STZ-icv) treatment and the transgenic Tg2576 mouse model of familial Alzheimer’s disease. It has been repeatedly shown that oxidative stress plays a pathogenic role in both models. As expected, NRP confirmed that TAC was reduced in plasma of both models (**Fig 2A**, **Fig 2B**). We proceeded to test TAC of hippocampal samples. Here, we needed to control for the amount of protein in each sample, as the same volume (1 μl) of all samples was pipetted to the nitrocellulose membrane for more precise and convenient analysis. As simple ratiometric correction assumes a linear relationship between NRP and protein content, this was tested prior to analysis with sampling-unadjusted and sampling-corrected protein estimation techniques. Protein content correction was justified for samples that dispersed from the representative sample by less than 50% of its protein concentration value (**Supplement 9**), so adjusted hippocampal NRP values were calculated and visualized. There was no difference between Tg2576 mice and their respective controls, and STZ-icv treatment demonstrated a trend towards decreased TAC (**Fig 2C**, **Fig 2D**). In summary, NRP can be used to rapidly obtain reliable information on reductive potential from small volumes of biological samples. Due to its simplicity, the method is very robust and convenient for parallel analysis of a large number of samples. Moreover, the method is compatible with different tissue sample preparation procedures so it can be easily combined with other analytical techniques and implemented in experimental design. However, NRP can also be adapted for analysis of anatomical distribution of reductive potential in solid tissue samples, a feature that makes our method unique and different from all other standard methods used for TAC evaluation. The HistoNRP method overview is demonstrated (**Fig 2E**) and detailed step-by-step protocols are available in **Supplement 1**. In short, proteins from tissue sections are transferred and fixed onto a nitrocellulose membrane by passive diffusion slice print blotting for cryosections and enzymatic retrieval followed by heat-facilitated passive diffusion slice printing for paraffin-embedded tissues. After fixation of proteins with preserved anatomical relations onto the nitrocellulose membrane, standard NRP protocol is used to obtain anatomical distributions of reductive capacity based on MnO_2_ precipitation patterns. A representative HistoNRP nitrocellulose membrane after passive diffusion slice print blotting of cryosections is shown in **Fig 2F** (an adaptation of the HistoNRP for analysis of paraffin-embedded tissue is demonstrated in **Supplement 1** and discussed in detail in **Supplement 10**). Once digitalized, the membranes can be further analyzed to obtain detailed information on spatial distribution of reductive capacity. For demonstration of spatial reductive analysis we used a rat brain damaged by microdialysis probe placement (a detailed description of the experiment can be found in the **Supplement 8**). Reductive capacity distribution of the damaged side of the brain was compared to the contralateral signal in order to explore how trauma affected different anatomical regions. Fire lookup table-based 3D intensity surface plot of reductive potential capacity obtained with HistoNRP protocol is shown in **Fig 2G**. Intensities of ipsilateral and contralateral pixels obtained by sagittal mirroring were visualized by fire lookup table-based 3D intensity surface plot (**Fig 2H**, left) and line profiles of interest were defined and visualized in (**Fig 2H**, right). Visual representation of individual intensity profiles from (**Fig 2H**) is presented in **Fig 2I**. Furthermore, brain areas of interest were identified and their intensity histograms were compared to the ones obtained from the contralateral side. Density plots of analyzed areas were sorted by their distance from the trauma site and visualized in **Fig 2J**. Detailed explanation of the analysis protocol is available in **Supplement 11**.

**Fig. 2.**
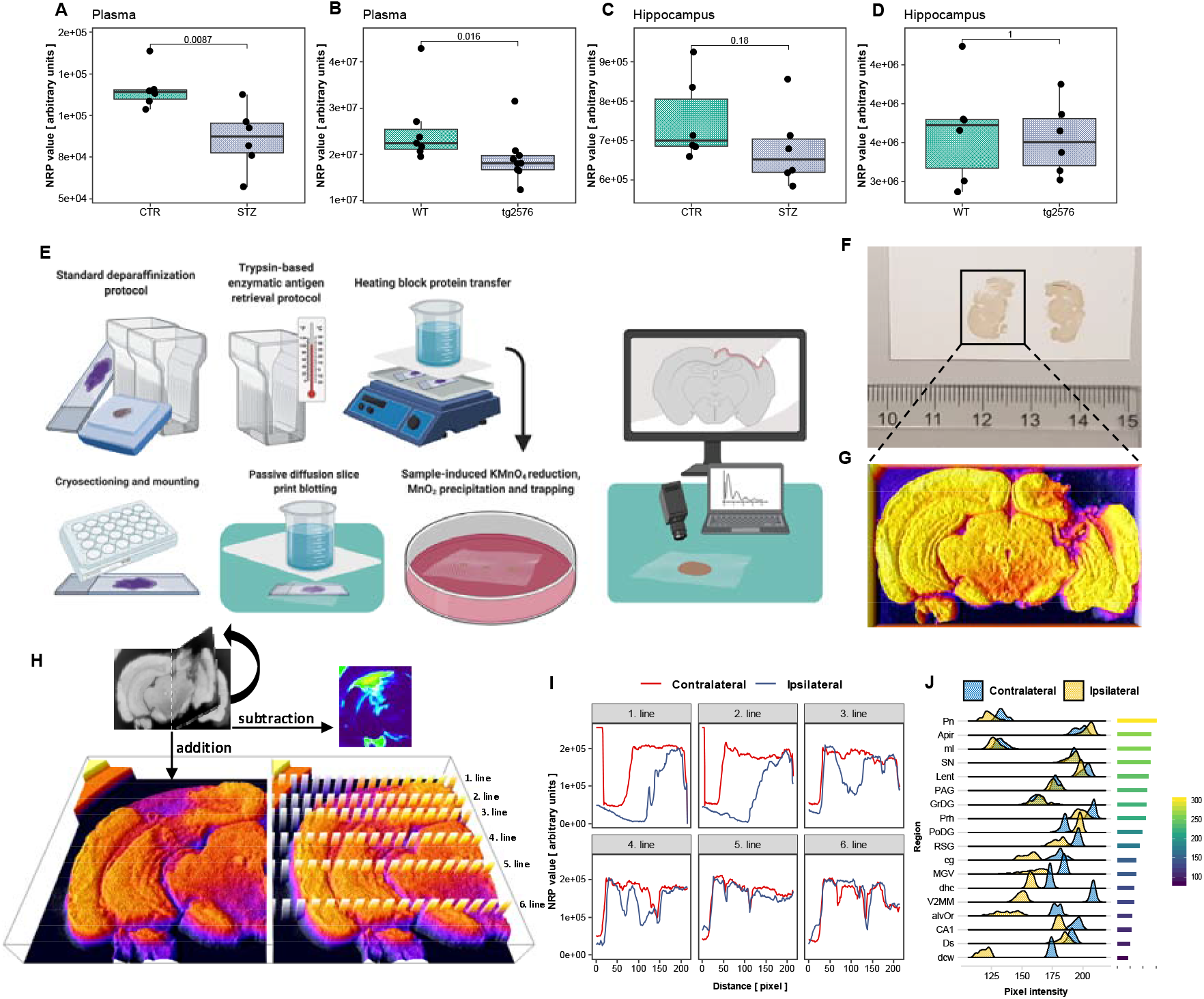
Demonstration of nitrocellulose redox permanganometry (NRP). **A**) Comparison of plasma reductive capacity of rats treated intracerebroventricularly with streptozotocin (STZ-icv) 1 month after the treatment (detailed description of the experiment is available in the **Supplement 8**). Plasma samples were placed on a nitrocellulose membrane in the volume of 1 μl. Once dry, the membrane was analyzed using the protocol described in the main text, digitalized, and integrated density of the signal was calculated for every sample in Fiji software using the gel analyzer tool. Groups were compared by Wilcoxon rank-sum test using R. **B**) The procedure described in (A) was repeated with plasma samples of transgenic animals (Tg2576) and their wild type controls (detailed description of the experiment is available in the **Supplement 8**). **C**) Hippocampal reductive potential was measured in tissues of the same animals from (A). In short, hippocampal homogenates were placed on a nitrocellulose membrane and the same protocol was followed as described in (A). After integrated density values were calculated, protein content correction was done by means of ratiometric normalization (the analysis of ratiometric protein correction approach is available in the **Supplement 9**). **D**) Hippocampal reductive potential was measured in tissues of the same animals described in (B). In short, hippocampal homogenates were placed on a nitrocellulose membrane and the same protocol was followed as described in (A). After integrated density values were calculated, protein content correction was done by means of ratiometric normalization. **E**) Schematic step-by-step overview of the NRP protocol adapted for analysis of histological specimens (HistoNRP). Briefly, formalin-fixed paraffin-embedded tissue (FFPE) is fixed onto a membrane by heat-facilitated passive diffusion slice printing, and cryosections are transferred onto nitrocellulose by means of passive diffusion slice print blotting. FFPE tissue is treated by standard deparaffinization protocol, followed by trypsin-based enzymatic antigen retrieval protocol, as commonly used for immunohistochemistry. After enzymatic digestion, the tissue is placed on a glass plate on a heating block. The nitrocellulose membrane is placed on tissue sections wetted with phosphate buffered saline (PBS) and covered with three wet filter papers, and the entire setup is covered with Parafilm^®^ and an additional glass plate. The heating block is set at 60°C and an 800 g weight is placed on top of the upper glass plate. The transfer is stopped after 8 hours (**Supplement 1**, **Supplement 10**), the membrane is wetted with PBS, carefully removed from the microscope slides and dried out. Once dry, the membrane is treated with the standard NRP protocol described earlier. For cryosections, once the tissue is mounted on the microscope slide and air-dried at 37°C, the slides are placed on a clean laboratory surface and wetted with PBS. A clean sheet of nitrocellulose membrane is placed on top of the slides and covered with three filter papers pre-wetted with PBS. The filter papers are covered with a glass plate and an 800 g weight is placed on top of the plate and left over night. The next day, the weight and filter papers are removed, and the membrane is wetted with PBS, carefully removed from the slides and left to air-dry. Once dry, the membrane is treated with the standard NRP protocol described earlier. **F**) An example of a nitrocellulose membrane of brain cryosections obtained by HistoNRP (a detailed description of the experiment is available in the **Supplement 8**). **G**) A brain cryosection used for demonstration of anatomical distribution of reductive capacity. The HistoNRP membrane was digitalized and analyzed in Fiji software. First, the image was turned to grayscale and inverted so pixel intensity represents reductive capacity. The fire lookup table-based 3D intensity surface plot is pictured. The section was chosen for demonstration due to evident brain damage caused by microdialysis probe placement. **H**) In order to analyze the reductive potential difference on the ipsilateral and contralateral side of the brain, the original photomicrograph was divided across the medial line. Sagittal mirroring of the damaged (ipsilateral) to control (contralateral) side was performed and intensity analysis was done by pixel addition and subtraction. The summation image is presented as a fire lookup table-based 3D intensity surface plot (bottom left). Lines of interest for analysis of dorso-ventral reductive capacity distribution were selected and visualized (bottom right). **I**) Pixel intensity profile plots of ipsilateral and contralateral contribution values were calculated and presented for each line of interest in dorso-ventral orientation. **J**) We identified 18 brain areas of interest, calculated pixel intensity histograms for all areas on the ipsilateral and contralateral side of the brain and measured pixel distances from ipsilateral areas to the trauma site. Ipsilateral and contralateral pixel value density plots of all areas were sorted by their distance to the trauma site (visualized in the legend on the right) to examine whether areas closest to the trauma site were also the ones with the greatest oxidative increment. **Pn** – paranigral nucleus; **Apir** – amygdalopiriform transition area; **ml** – medial lemniscus; **SN** – substantia nigra; **Lent** – lateral entorhinal cortex; **PAG** – periaqueductal gray area; **GrDG** – granular dentate gyrus; **Prh** – perirhinal cortex; **PoDG** – polymorph layer of the dentate gyrus; **RSG** – retrosplenial granular cortex; **cg** – cingulum; **MGV** – medial geniculate nucleus, ventral part; **dhc** – dorsal hippocampal commissure; **V2MM** – secondary visual cortex, mediomedial area; **alvOr** – alveus of the hippocampus and oriend layer of the hippocampus; **CA1** – field CA1 of the hippocampus; **Ds** – dorsal subiculum; **dcw** – deep cerebral white matter.

In conclusion, our results demonstrate that NRP is a rapid, cost-effective and simple method that can be used for high-throughput analysis of reductive potential. Moreover, its extension, HistoNRP provides invaluable information on tissue distribution of reductive capacity in great detail and can be used in concordance with other methods for thorough understanding of oxidative stress.

## Supporting information

Supplement 1 - A step-by-step Nitrocellulose redox permanganometry (NRP) and HistoNRP protocols

Supplement 2 - Nitrocellulose redox permanganometry (NRP) linearity, precision and accuracy validation tests

Supplement 3 - Nitrocellulose redox permanganometry (NRP) ascorbic acid heating time and temperature response validation experiments

Supplement 4 - Nitrocellulose redox permanganometry (NRP) membrane stability analysis

Supplement 5 - Comparison of Nitrocellulose redox permanganometry (NRP) digitalization techniques

Supplement 6 - An explanation of the computational analysis of Nitrocellulose redox permanganometry (NRP)

Supplement 7 - Additional replications of ORP-NRP experiments from Fig 1E and Fig 1F

Supplement 8 - A detailed explanation of the origin of animal tissue used in the Nitrocellulose redox permanganometry (NRP) proof of concept experimen

Supplement 9 - Nitrocellulose redox permanganometry (NRP) protein concentration-based correction analysis

Supplement 10 - HistoNRP adaptation for the analysis of formalin-fixed paraffin-embedded (FFPE) tissue sections

Supplement 11 - A detailed explanation of the HistoNRP demonstration analysis illustrated in the Fig 2 of the Main text

## Authors contributions

**JH** conceptualized the method, designed experimental protocols and validation tests, and conducted all NRP experiments. **JH**, **IK** and **DV** conducted data analysis and visualization. **JH**, **IK**, **ABP** and **DV** wrote the manuscript. **JH**, **ABP**, **AK** and **JOB** conducted animal experiments from which tissue samples were taken for proof of concept analyses. **MSP** is head of the Laboratory for Molecular Neuropharmacology where all experiments were conducted, principal investigator of the Croatian Science Foundation funded project (IP-2018-01-8938) and research group leader in the Scientific Centre of Excellence for Basic, Clinical and Translational neuroscience (GA KK01.1.1.01.0007) and supervisor and mentor of the authors. **PR** provided valuable comments on the manuscript and the method as a renowned expert in the field of neurochemistry and a close collaborator of the group. All authors approved the manuscript and provided comments during the writing process.

## Ethics committee approval

All experiments were conducted in concordance with the highest standard of animal welfare. Only certified personnel handled animals. Animal procedures were carried out at the University of Zagreb Medical School (Zagreb, Croatia) and were in compliance with current institutional, national and international (The Animal Protection Act, NN 102/17, 32/19, NN 125/13, 14/14, 92/14, 32/19), and international (Directive 2010/63/EU) guidelines governing the use of experimental animals. The experiments were approved by the national regulatory body responsible for issuing ethical approvals, the Croatian Ministry of Agriculture and by the Ethical Committee of the University of Zagreb School of Medicine.

