## Supplement 1 - A step-by-step Nitrocellulose redox permanganometry (NRP) and HistoNRP protocols for "Nitrocellulose redox permanganometry: a simple method for reductive capacity assessment"

### A step-by-step Nitrocellulose Redox Permanganometry (NRP) protocol for liquid samples

#### Materials and reagents

- A piece of nitrocellulose membrane
- A small volume pipette
- Pipette tips
- $KMnO_4$
- $ddH_2O$
- Tweezers
- A Petri dish
- Glass beaker
- A stirring magnet
- Magnetic stirrer

#### NRP Protocol

1. **Prepare the $KMnO_4$ working solution by dissolving 0.2 g of solid $KMnO_4$ crystals in 20 mL of $ddH_2O$ (place the solution with a magnet on the magnetic stirrer until all crystals dissolve).**

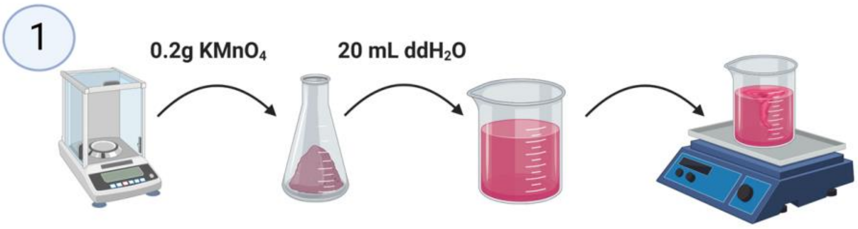


2. **Remove the protective cover from one side of the nitrocellulose membrane and place the membrane on a clean laboratory surface (leave the protective cover on the side of the membrane facing the laboratory surface).**

3. **Vortex samples.**

4. **Pipette 1 μL of the sample onto the nitrocellulose membrane, repeat this step with every sample you want to analyze.**

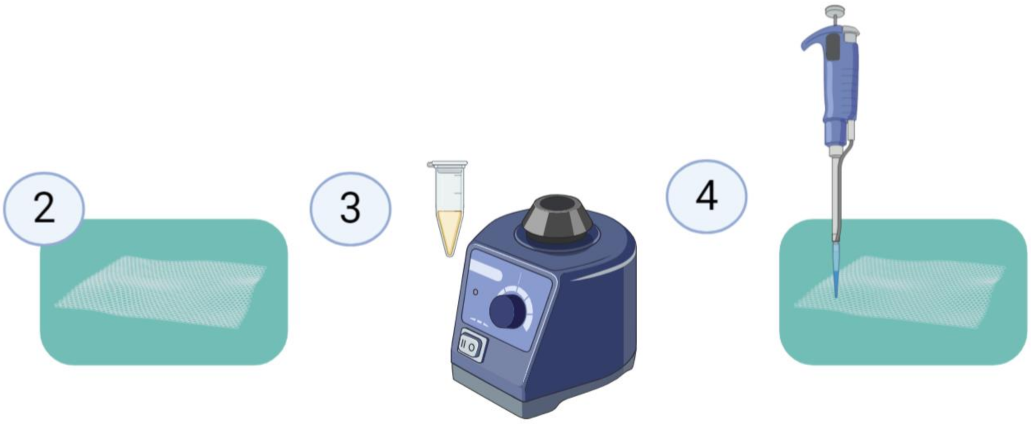


**Caution: Leave enough space between your samples.**

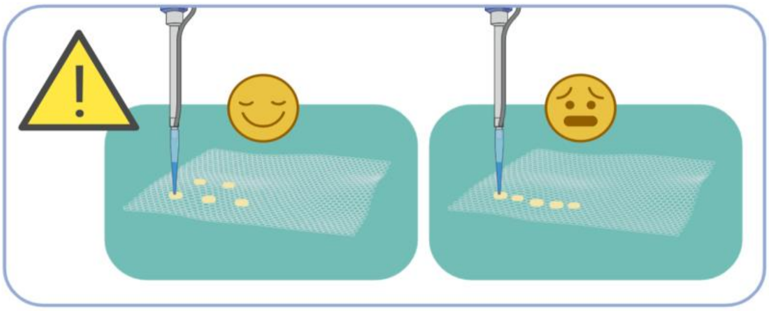

5. **Leave the nitrocellulose membrane to dry out.**

6. **Once dry, pick up the nitrocellulose membrane with tweezers and place it in the $KMnO_4$ working solution.**

7. **Wait for 30 seconds.**

8. **Remove the nitrocellulose membrane from the $KMnO_4$ working solution with tweezers and place it under running $ddH_2O$ to terminate the reaction and increase the contrast.**

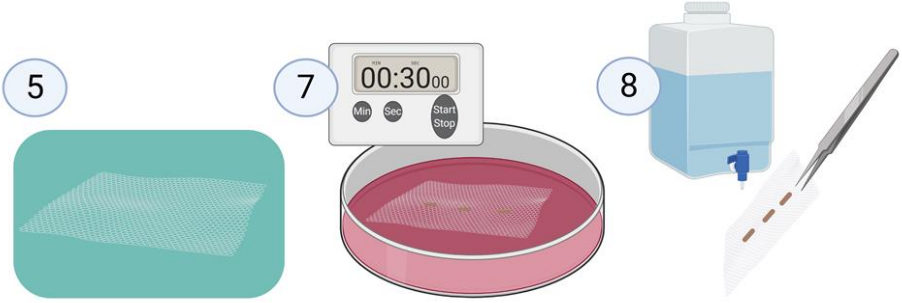


9. **Leave the nitrocellulose membrane to dry out.**

10. **Digitalize the membrane (with a cellphone, camera or an office scanner).**

11. **Import the photo into the Fiji (Fiji is just ImageJ) software.**

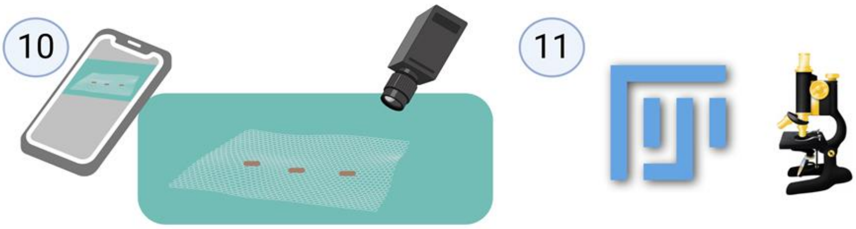


12. **Select the region of interest in Fiji by using the "Rectangle selection" tool.**

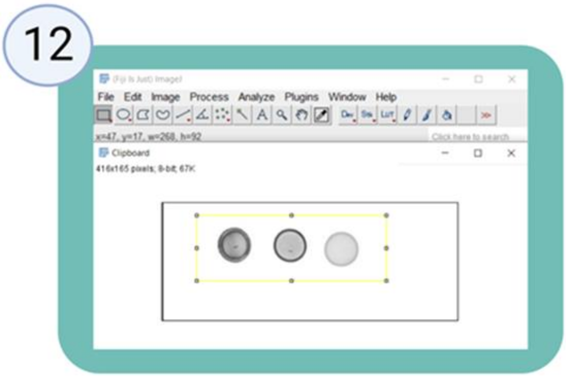

13. **Perform the Gel Analyzer plugin “Select First Lane” function (CTRL+1).**

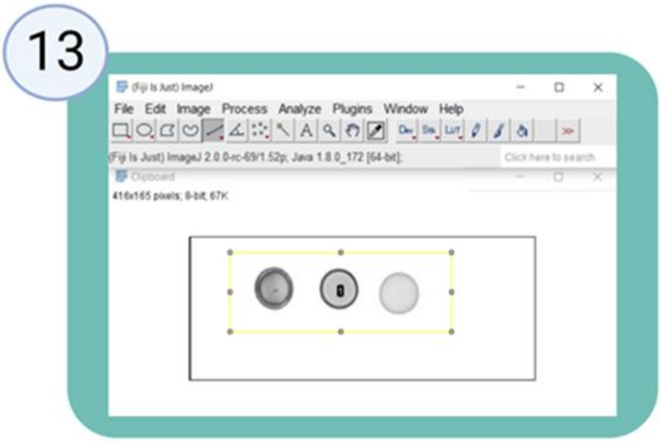


14. **Perform the Gel Analyzer plugin “Plot Lanes” function (CTRL+3).**

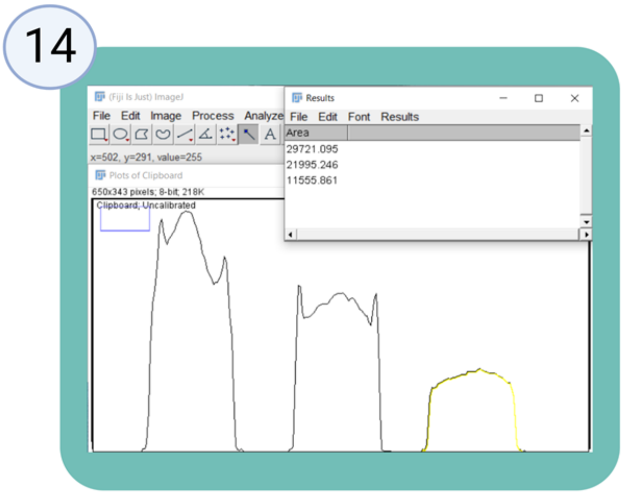


15. **Export values from the FIji “Results” tab, and proceed to statistical analysis and visualization (greater values represent higher reductive (antioxidant) capacity).**

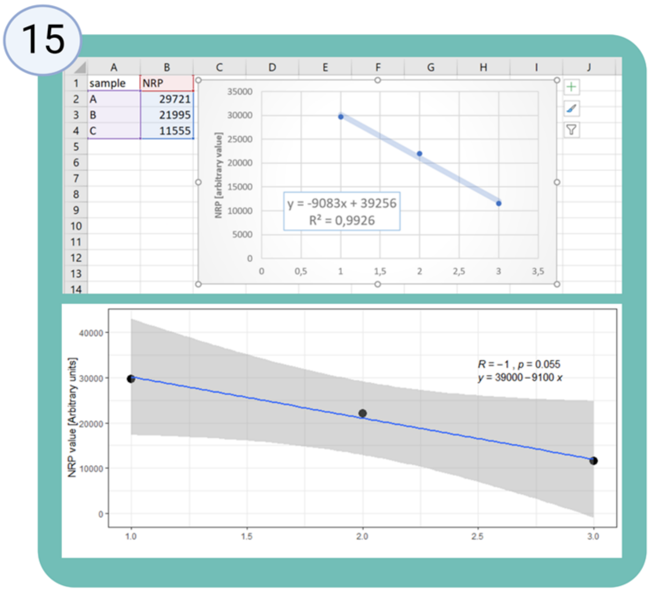


#### A step-by-step HistoNRP protocol for cryosections

#### Materials and reagents

- A piece of nitrocellulose membrane
- Filter papers
- A pipette
- $KMnO_4$
- dd$H_2O$
- Phosphate buffered saline (1xPBS)
- Tweezers
- A Petri dish
- Glass beaker
- A glass plate
- A stirring magnet
- Magnetic stirrer

### HistoNRP protocol for cryosections

1. Prepare the $KMnO_4$ working solution by dissolving 0.2 g of solid $KMnO_4$ crystals in 20 mL of $ddH_2O$ (place the solution with a magnet on the magnetic stirrer until all crystals dissolve).

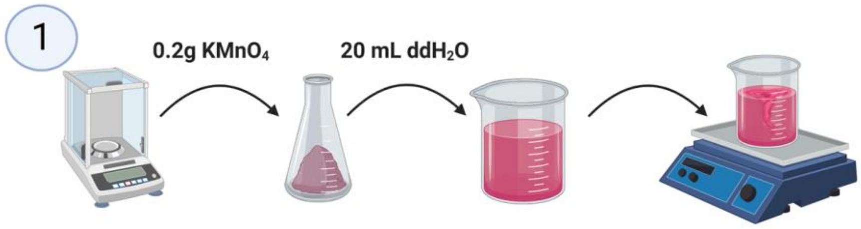


2. Remove the frozen tissue samples from the freezer and use a cryostat to make cryosections of the tissue of interest. Place the sections directly onto the glass slides. Air dry the samples at 37°C.

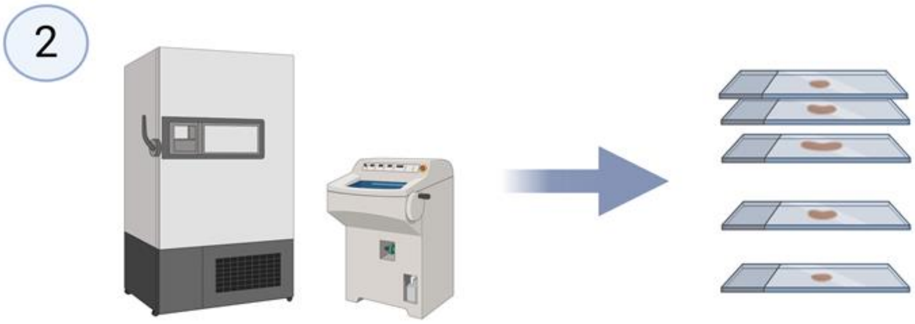


3. Place the slides onto a clean laboratory surface and wet the tissue samples with PBS.
4. Place a piece of nitrocellulose membrane onto the wetted slide.
5. Place 3 filter papers on top of the membrane.
6. Wet the filter papers with PBS.

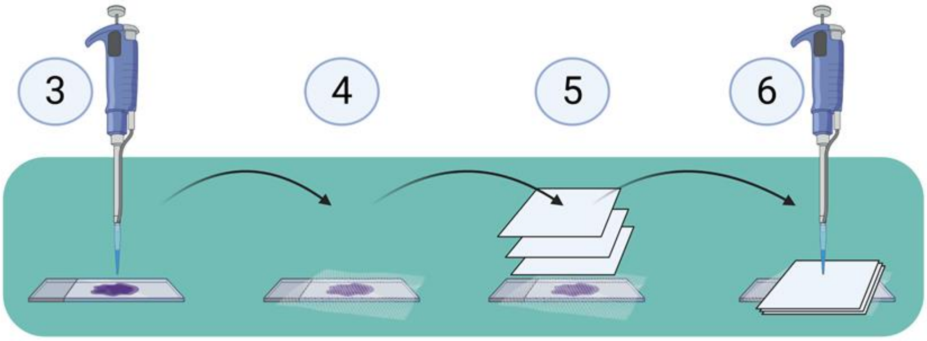


7. Cover everything with a glass plate for even distribution of the weight and place a full beaker onto the glass plate to apply appropriate pressure (the optimal pressure should be determined by the end user; in our case it was 31.384 mmHg achieved by placing a

**beaker with 800 g of water on the glass plate). Leave the sample proteins to passively diffuse onto the membrane over night at room temperature.**

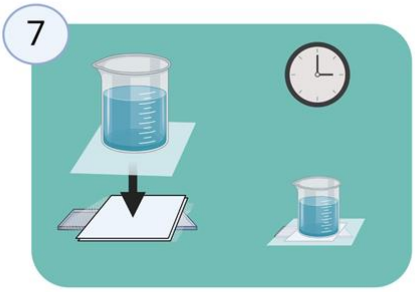


8. **Remove the beaker, glass plate, and 2 filter papers carefully. Wet the last filter paper with PBS and remove it carefully with tweezers. Add additional PBS and remove the membrane from the slide. Leave the membrane with printed proteins to dry out.**

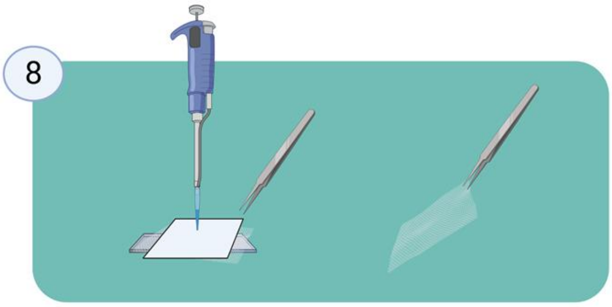


9. **Once dry, pick up the nitrocellulose membrane with tweezers and place it in the $KMnO_4$ working solution for 30 seconds.**

10. **Remove the nitrocellulose membrane from the $KMnO_4$ working solution with tweezers and place it under running dd$H_2O$ to terminate the reaction and increase the contrast.**

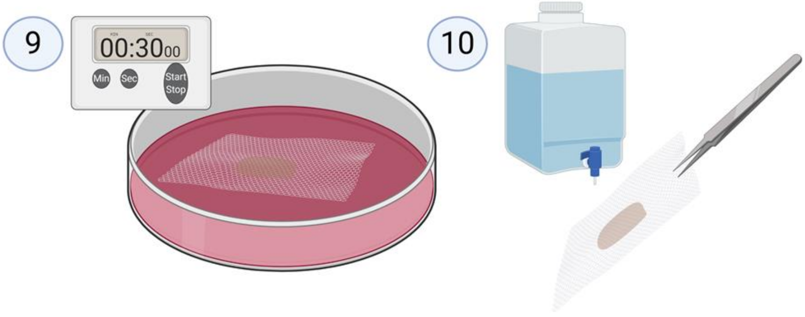


11. **Leave the nitrocellulose membrane to dry out and digitalize the membrane (with a cellphone, camera or an office scanner).**
12. **Import the photo into the Fiji (Fiji is just ImageJ) software.**

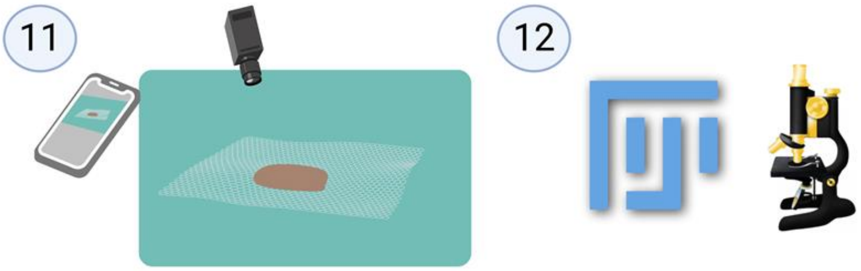

13. **Prepare the image for intensity analysis by 8-bit (Image>Type>8-bit) processing and appropriate inversion (Edit>Invert; "CTRL+shift+I").**

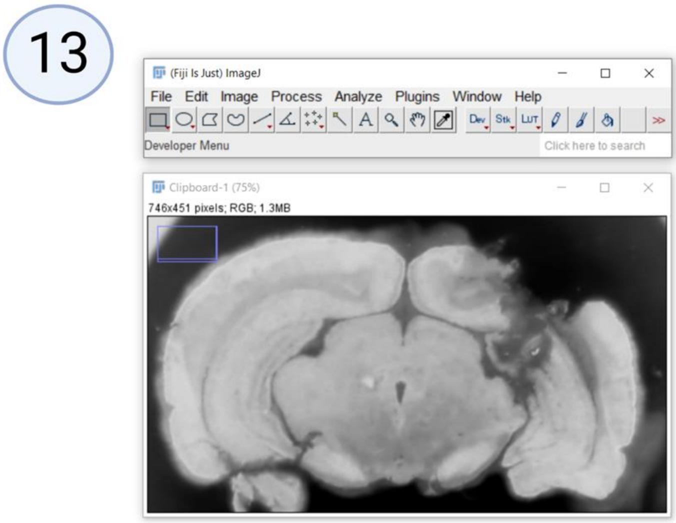


14. **Select the area of interest using the selection tool and measure pixel intensities by using the Fiji "Histogram" function (Analyze>Histogram; "CTRL+H").**

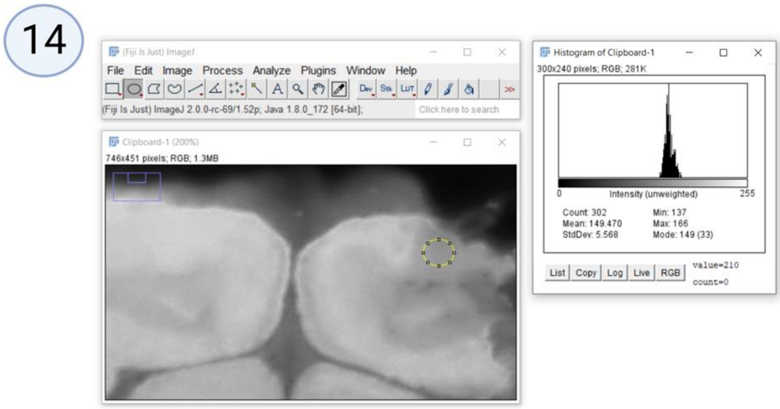


15. **Use the "List" option to export values from Fiji for further analysis.**

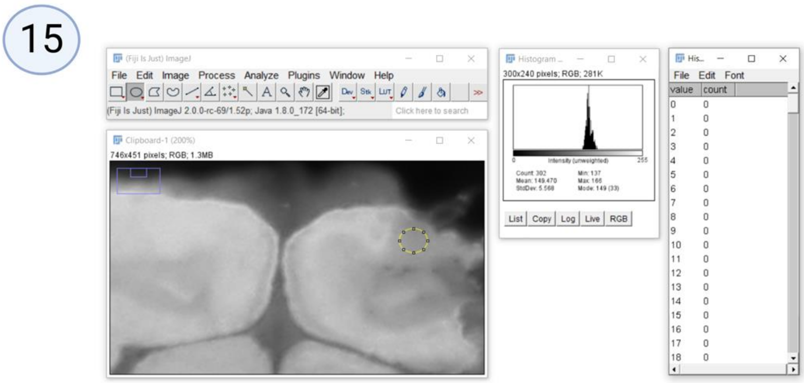


16. **An example of pixel intensity comparisons from the ipsilateral side following microdialysis probe-induced damage and the contralateral side of the brain used as a reference. A comparative analysis of 18 brain areas analyzed by density plot comparison**

**done in R software environment for statistical computing is further explained in the Main text.**

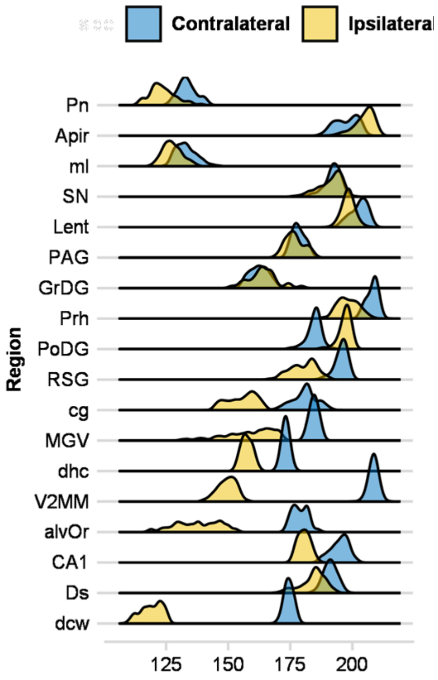

### A step-by-step HistoNRP protocol for formalin-fixed paraffin-embedded (FFPE) tissue samples

#### Materials and reagents

- A piece of nitrocellulose membrane
- Filter papers
- A pipette
- $KMnO_4$
- $ddH_2O$
- Phosphate buffered saline (1xPBS)
- Tweezers
- A Petri dish
- Glass beaker
- Two glass plates
- A stirring magnet
- Magnetic stirrer
- Laboratory heater
- Parafilm
- Equipment for FFPE tissue deparaffinization (Xylene, EtOH)

#### HistoNRP protocol for FFPE samples

1. **Prepare the $KMnO_4$ working solution by dissolving 0.2 g of solid $KMnO_4$ crystals in 20 mL of $ddH_2O$ (place the solution with a magnet on the magnetic stirrer until all crystals dissolve).**

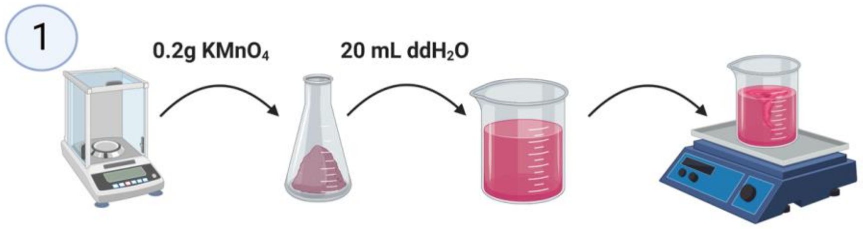


2. **Cut the FFPE tissue on the microtome and mount the sections on histological slides.**

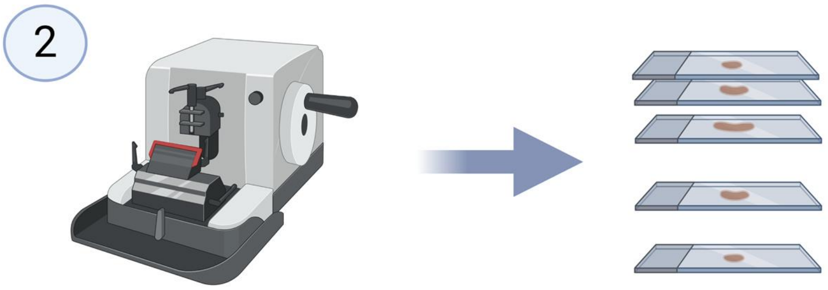


3. **Deparaffinize the FFPE slides using a standard procedure (eg. 3x5 min Xylene/ 2x5 min 100% EtOH/ 2x5 min 95% EtOH/ 2x5 min 70% EtOH/ 2x5 min 50% EtOH) and place the slides in PBS (2x5 min).**

4. **Place the slides in a warm (37°C) antigen retrieval solution (0.05% Trypsin, 0.1% CaCl in $ddH_2O$; pH 7.8) for 45 minutes.**

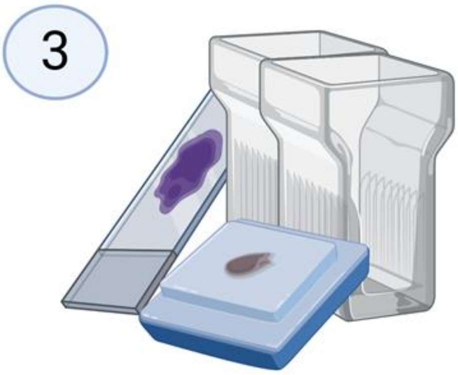


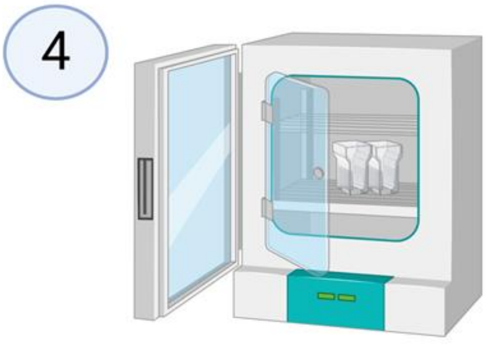


5. **Place the slides in PBS (2x5 min) at room temperature to acclimate and prepare the heat-facilitated passive diffusion slice printing setup. The beaker should provide approximately the same pressure as in the protocol for cryosections (31.384 mmHg).**

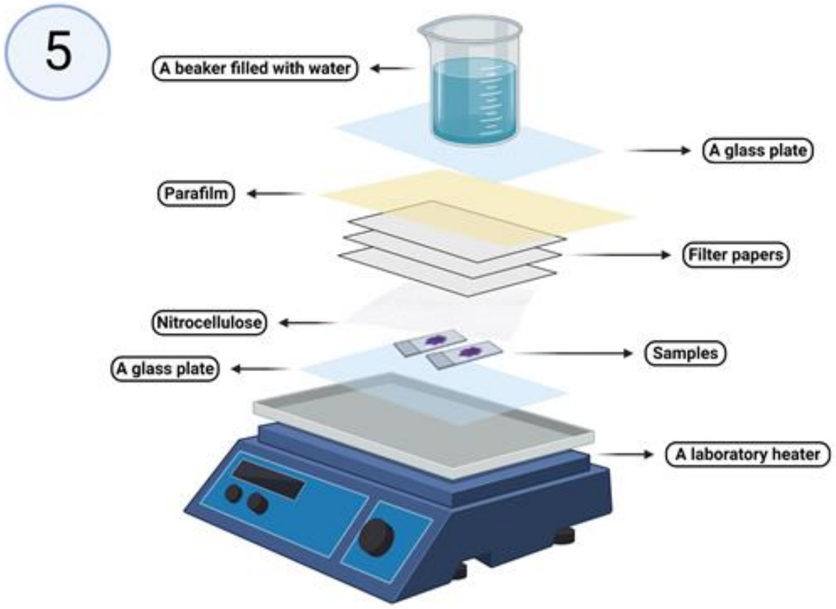


6. **Turn on the heater and press the parafilm towards the lower glass plate to ensure optimal isolation and humidity during the transfer protocol.**

7. **The transfer protocol should be modified by the end user; in our case the optimal transfer temperature for rat brain slices was 60°C and the optimal time was determined to be 8 hours.**

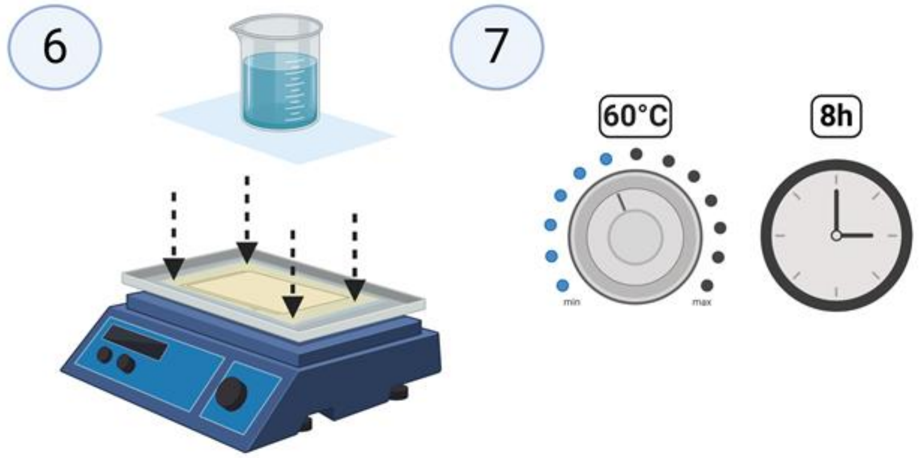


8. **After the heat-facilitated passive diffusion slice printing protocol is over, the beaker, upper glass plate, parafilm and 3 filter papers should be removed, and the membrane should be wetted with PBS, carefully removed with tweezers and left to dry.**

9. **Once dry, pick up the nitrocellulose membrane with tweezers and place it in the $KMnO_4$ working solution for 30 seconds.**

10. **Remove the nitrocellulose membrane from the $KMnO_4$ working solution with tweezers and place it under running $ddH_2O$ to terminate the reaction and increase the contrast.**

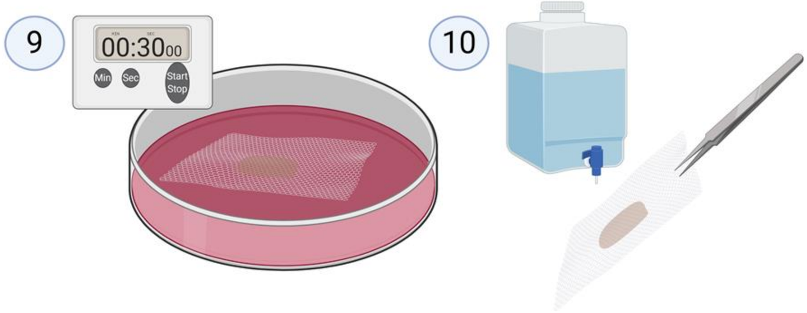


11. **Leave the nitrocellulose membrane to dry out and digitalize the membrane (with a cellphone, camera or an office scanner).**

12. **Import the photo into the Fiji (Fiji is just ImageJ) software.**

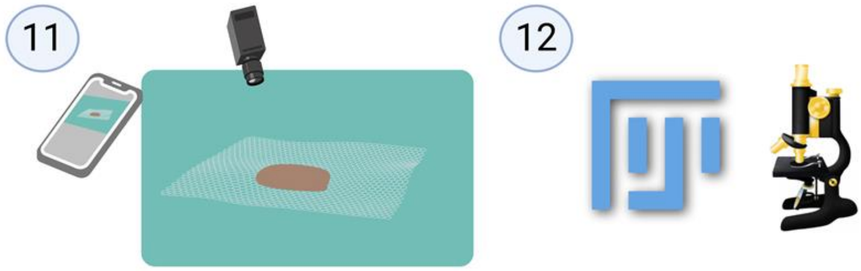


13. **Prepare the image for intensity analysis by 8-bit (Image>Type>8-bit) processing and appropriate inversion (Edit>Invert; "CTRL+shift+I").**

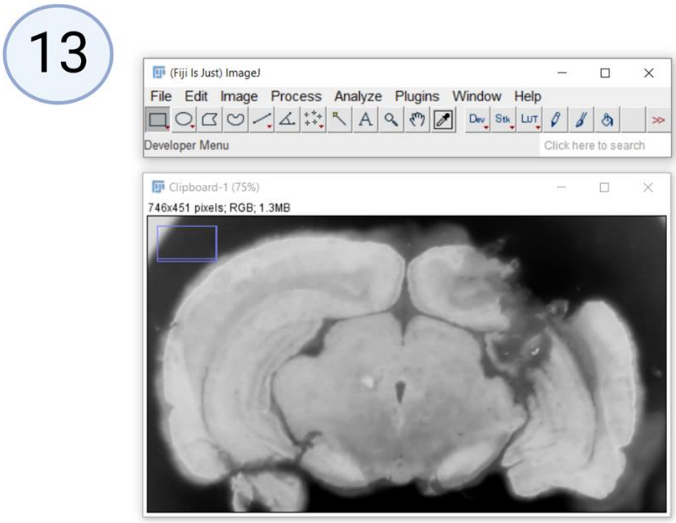


14. **Select area of interest using the selection tool and measure pixel intensities by using the Fiji "Histogram" function (Analyze>Histogram; "CTRL+H").**

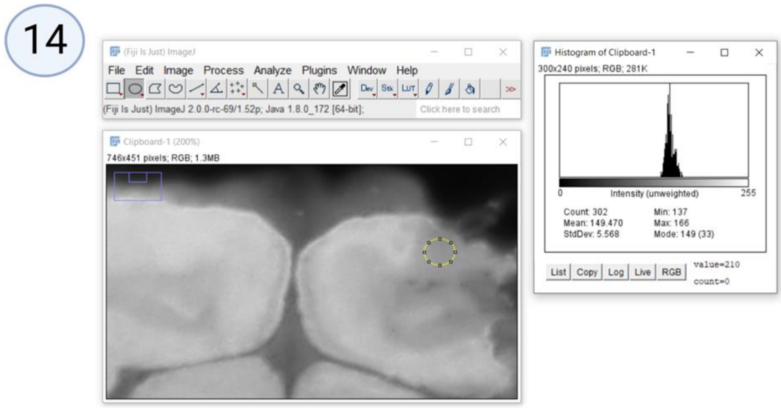


15. **Use the "List" option to export values from Fiji for further analysis.**

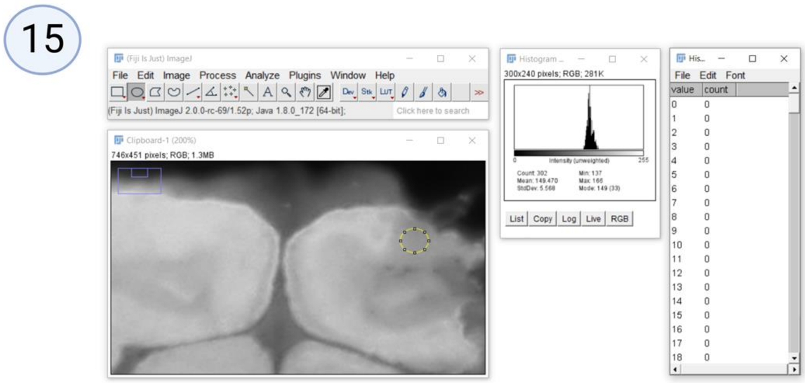


16. **An example of pixel intensity comparisons from the ipsilateral side following microdialysis probe-induced damage and the contralateral side of the brain used as a reference. A comparative analysis of 18 brain areas analyzed by density plot comparison done in R software environment for statistical computing is further explained in the Main text.**

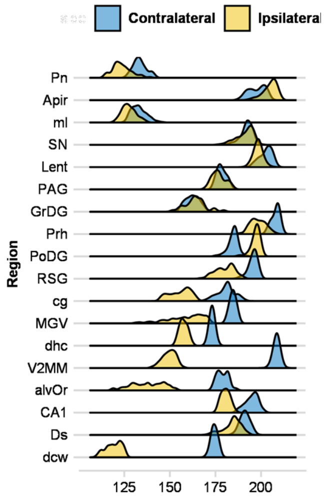
