## Supplement 2 - Nitrocellulose redox permanganometry (NRP) linearity, precision and accuracy validation tests for "Nitrocellulose redox permanganometry: a simple method for reductive capacity assessment"

In order to test linearity, accuracy and precision of nitrocellulose redox permanganometry (NRP), we prepared 10 stock solutions of a standard reductive agent, sodium thiosulfate ($Na_2S_2O_3$), often used for validation of the total antioxidant capacity method. Ten nominal concentrations between 0.1M – 0.01M were prepared independently by weighing out an appropriate amount of $Na_2S_2O_3$ and dissolving them in double-distilled water (dd$H_2O$) with predetermined purity of 0.055 µS/cm. Samples were placed at 4°C until everything was ready for NRP.

The European Medicines Agency Bioanalytical Method Validation Guidelines (EMA-BMVG)[1] suggest a minimum of 6 samples to be used for generation of the calibration curve (CC). Here, we used 10 nominal concentrations. Moreover, EMA-BMVG suggest each calibration standard can be analyzed in replicates. Here, 5 technical replicates per concentration were used. It is suggested that a relationship between analyte concentration and the response of the instrument should be reported, and considering that our results had a good linear fit, we reported a standard linear regression equation (y=2100+1.4e6x; $R^2$=0.99; p=1.3e-49) (**Main text Fig 1B**).

Accuracy was obtained for all ten nominal concentration points following the within-run-accuracy determination principles. EMA-BMVG suggest a minimum of 5 samples per level at a minimum of 4 concentration levels which are covering the CC range by the following principle: quality control (QC) samples have to cover the whole concentration curve and include samples reflecting low concentrations (low QC; below 30% of the CC range), medium concentrations (medium QC; around 30-50% of the CC range) and high concentrations (high QC; above 75% of the CC range). Here, we tested within-run accuracy with 5 samples per level at 10 concentration points. More precisely, EMA-BMVG suggest low QC to be tested at approximately 3 times the concentration of the lower limit of quantification (LLOQ), the lowest concentration value of analyte in a sample which can be quantified reliably. However, as most of the (biological) samples preliminarily tested fell onto the concentration curve generated by analysis of 0.01M - 0.1M $Na_2S_2O_3$, we didn't formally extend our concentration curve looking for the true potential LLOQ concentration point and following the EMA-BMVG principles our LLOQ point for the within-run $Na_2S_2O_3$ validation run probably could have been shifted towards much lower values. For example, for within-run accuracy, 20% deviation from the nominal values is considered acceptable for the LLOQ point. In our case, 0.01 $Na_2S_2O_3$ taken as LLOQ deviated from nominal values only by 9.47% (**Main text Fig 1B**). Within-run accuracy principles suggest mean concentrations to be within 15% of the nominal values for the low, medium and high QC, and within 20% for the LLOQ. Our $Na_2S_2O_3$ QC samples all had a coefficient of variation (CV) below 10%, and all except 0.01M $Na_2S_2O_3$ had CV below 4% (**Table S2-1**). As $Na_2S_2O_3$ validation run CC was calculated from the average integrated density of every QC data point, only CV was used as a validation parameter because mean group values didn't deviate from the CC values.

As explained in the main text, the whole validation protocol was repeated with another standard reductive agent, ascorbic acid (AA), to ensure that the test truly measured reductive capacity rather than some other property of $Na_2S_2O_3$. Here, we also decided to test a range of 10 nominal concentrations between 0.1M – 0.01M, although we expected different absolute reductive values. The reason for this was to simply demonstrate two qualities in the same run - the fact that the test really reflects reductive capacity of the tested samples, and to show its robustness. In this run the results obtained from the AA NRP test also had a good linear fit so a standard linear regression equation was reported (y=2700+5.9e5x; $R^2$=0.94; p=6.6e-33) (**Main text Fig 1C**). All validation tests were done following the same principles as described for $Na_2S_2O_3$ validation. Precision of the test was determined by calculating CV for all 10 samples (**Table S2-2**). As expected, CC for AA was much closer to LLOQ of the NRP. For example, 0.01 M AA had a CV of 21.27%, 1.27% above the recommended LLOQ range. Consequently, for samples with reductive capacity equal or lower than 0.01M AA in this experiment, NRP protocol would have to be adjusted to obtain reliable information.

**Table S2-1.** Raw data used for calculation of the $Na_2S_2O_3$ coefficient of variation in the NRP validation test

| Sodium thiosulfate concentration [M] | Raw NRP values [arbitrary units] | CV [%] |
|---|---|---|
| 0.1 | 125412 | 3.765271544 |
| 0.1 | 129138 | |
| 0.1 | 131512 | |
| 0.1 | 138604 | |
| 0.1 | 133835 | |
| 0.09 | 122305 | 0.895188076 |
| 0.09 | 121010 | |
| 0.09 | 121694 | |
| 0.09 | 121434 | |
| 0.09 | 119424 | |
| 0.08 | 110419 | 2.763652803 |
| 0.08 | 112968 | |
| 0.08 | 114326 | |
| 0.08 | 117297 | |
| 0.08 | 118158 | |
| 0.07 | 106306 | 2.471816343 |
| 0.07 | 103381 | |
| 0.07 | 101882 | |
| 0.07 | 101888 | |
| 0.07 | 99383 | |
| 0.06 | 83345 | 2.724647765 |
| 0.06 | 84941 | |
| 0.06 | 86134 | |
| 0.06 | 87831 | |
| 0.06 | 89329 | |

| | | |
|---|---|---|
| 0.05 | 76423 | 1.830637261 |
| 0.05 | 77698 | |
| 0.05 | 77378 | |
| 0.05 | 77823 | |
| 0.05 | 74438 | |
| 0.04 | 51815 | 2.643992359 |
| 0.04 | 50511 | |
| 0.04 | 53308 | |
| 0.04 | 54021 | |
| 0.04 | 53084 | |
| 0.03 | 42946 | 3.068341054 |
| 0.03 | 43876 | |
| 0.03 | 45913 | |
| 0.03 | 44383 | |
| 0.03 | 42640 | |
| 0.03 | 42282 | |
| 0.02 | 27454 | 3.300032665 |
| 0.02 | 27303 | |
| 0.02 | 26776 | |
| 0.02 | 28740 | |
| 0.02 | 28628 | |
| 0.02 | 28990 | |
| 0.01 | 14150 | 9.473625281 |
| 0.01 | 14780 | |
| 0.01 | 14083 | |
| 0.01 | 12195 | |
| 0.01 | 11952 | |
| 0.01 | 9103 | |

**Table S2-2.** Raw data used for calculation of the ascorbic acid coefficient of variation in the NRP validation test

| Ascorbic acid concentration [M] | Raw NRP values [arbitrary units] | CV [%] |
|---|---|---|
| 0.1 | 55036 | 2.327915513 |
| 0.1 | 58096 | |
| 0.1 | 56067 | |
| 0.1 | 54797 | |
| 0.1 | 56203 | |
| 0.09 | 54906 | 2.374952486 |
| 0.09 | 56918 | |
| 0.09 | 57860 | |
| 0.09 | 58323 | |
| 0.09 | 57784 | |
| 0.08 | 44425 | 3.005992443 |
| 0.08 | 47286 | |
| 0.08 | 47712 | |
| 0.08 | 47574 | |
| 0.08 | 47660 | |
| 0.07 | 44199 | 3.532410204 |
| 0.07 | 47128 | |
| 0.07 | 47707 | |
| 0.07 | 48025 | |
| 0.07 | 49138 | |
| 0.07 | 46719 | |
| 0.06 | 37750 | 2.315678687 |
| 0.06 | 39550 | |
| 0.06 | 39183 | |
| 0.06 | 39453 | |
| 0.06 | 37802 | |
| 0.05 | 33809 | 6.171647379 |
| 0.05 | 36066 | |
| 0.05 | 36669 | |
| 0.05 | 39103 | |
| 0.05 | 39293 | |
| 0.04 | 27346 | 4.44893066 |
| 0.04 | 30021 | |
| 0.04 | 30672 | |
| 0.04 | 30299 | |
| 0.04 | 29860 | |
| 0.03 | 22104 | 7.877937712 |
| 0.03 | 24360 | |
| 0.03 | 25784 | |
| 0.03 | 27988 | |
| 0.03 | 26430 | |

| | | |
|---|---|---|
| 0.03 | 25654 | |
| 0.02 | 9572 | 12.91127875 |
| 0.02 | 10793 | |
| 0.02 | 12446 | |
| 0.02 | 13113 | |
| 0.02 | 10495 | |
| 0.01 | 1809 | 21.27127437 |
| 0.01 | 1978 | |
| 0.01 | 2658 | |
| 0.01 | 2339 | |
| 0.01 | 2491 | |
| 0.01 | 3250 | |
