## Supplement 3 - Nitrocellulose redox permanganometry (NRP) ascorbic acid heating time and temperature response validation experiments for "Nitrocellulose redox permanganometry: a simple method for reductive capacity assessment"

We further tested the specificity of NRP towards reductive potential to exclude the possibility that the test is in fact reflecting some property related to the concentration of both $Na_2S_2O_3$ and ascorbic acid (AA). Usually, methods are validated only by concentration curve linearity and precision analysis; however, we believe that the principle of validation that relies exclusively on the concentration-signal relationship analysis is logically flawed as substance concentration bias is unavoidable. In order to overcome this problem, we designed experiments based on the idea of physical oxidation of isoconcentrated samples. Being a classic antioxidant with good NRP linearity and precision (**Main text Fig 1C, Supplement 2**), and known for its thermal sensitivity, we decided to use heat-induced decrement of AA reductive capacity as a proof-of-concept for isoconcentrated substance validation. Three experiments were performed. First, we independently prepared six solutions of 0.05M AA and aliquoted each sample into two Eppendorf tubes. One of two aliquots was placed at 4°C and the other in a heating block at 70°C. After 4 hours, both aliquots were left to acclimate to room temperature and then 1 µl of each solution was placed on a clean sheet of nitrocellulose membrane. Standard NRP protocol was applied and the results were presented in **Fig S3-1A** and **Main text Fig 1D**. As expected, reductive capacity was reduced upon heating ($p=0.0022$). We then moved on to further test this by analyzing the relationship between NRP signal reduction and AA heating time. We prepared 6 replicates of AA samples with three aliquots and placed them either at 4°C for 6.5 hours, at 4°C for 2.5 hours followed by 4 hours at 70°C, or at 70°C for 6.5 hours. Afterwards, all samples were left to acclimate to room temperature and then 1 µl of each solution was placed on a clean sheet of nitrocellulose membrane and standard NRP protocol was conducted. Time response of AA oxidation is presented in **Fig S3-1B**. At the temperature of 70°C most of the AA oxidation occurred in the first 4 hours. Prolongation of AA heating for 2.5 hours provided a small additional decrement of AA reductive potential. In addition, we performed a temperature response experiment to see whether heating at higher temperatures would exert a greater decrement of AA reductive capacity. Here, we independently prepared duplicate samples of 0.1M AA with 4 aliquots and placed the samples for 4 hours at either 4°C, 50°C, 70°C or 95°C. Afterwards, all samples were left to acclimate to room temperature and then 1 µl of each solution was placed on a clean sheet of nitrocellulose membrane for NRP. The results are presented in **Fig S3-1C**. Higher temperature resulted in a greater decrease of NRP values. More specifically, 4 hours at 50°C in this experiment reduced AA NRP by approximately 25%, 4 hours at 70°C by 48%, and 4 hours at 95°C by 63%. In conclusion, time and temperature responses of isoconcentrated AA reductive capacity suggest that the results obtained by NRP reflect antioxidant capacity rather than some other concentration-dependent variable.

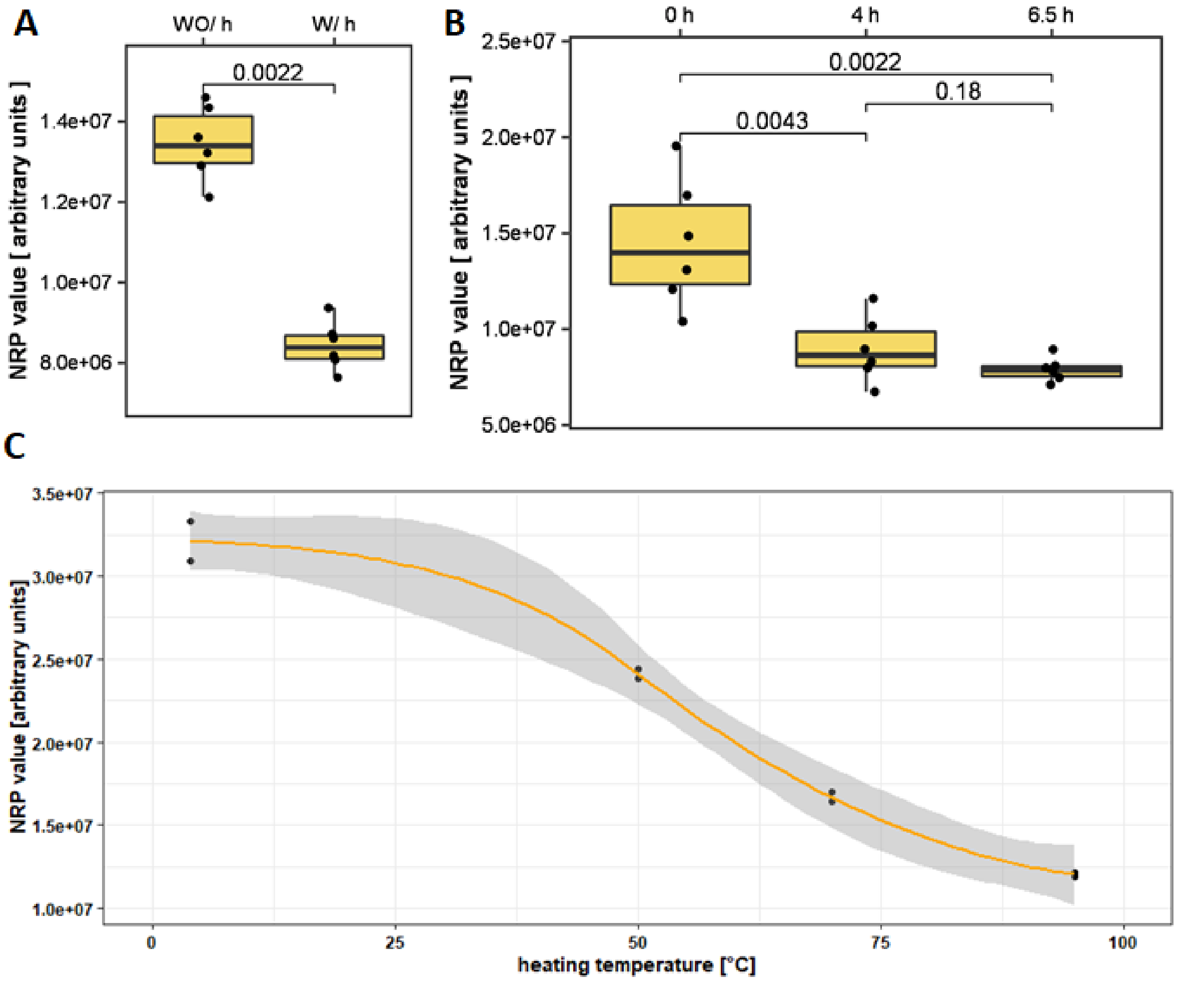


**Fig S3-1. Nitrocellulose redox permanganometry (NRP) ascorbic acid heating time and temperature response validation experiments. A)** Six independent replicates of 0.05M ascorbic acid (AA) in duplicate aliquots were heated at either 70°C for 4 hours or left at 4°C. After samples acclimated to room temperature, 1 µl of each solution was placed on a clean sheet of nitrocellulose membrane and analyzed by a standard NRP protocol. **B)** Six replicates of AA samples with three aliquots were placed at either 4°C for 6.5 hours, at 4°C for 2.5 hours followed by 4 hours at 70°C, or at 70°C for 6.5 hours. After heating, the samples were acclimated to room temperature and analyzed by NRP. **C)** Independently prepared duplicate samples of 0.1M AA with 4 aliquots were placed for 4 hours at either 4°C, 50°C, 70°C or 95°C. Samples were acclimated to room temperature and analyzed by NRP. NRP-nitrocellulose redox permanganometry; WO/ h-without heating; W/ h-with heating.
