## Supplement 4 - Nitrocellulose redox permanganometry (NRP) membrane stability analysis for "Nitrocellulose redox permanganometry: a simple method for reductive capacity assessment"

Nitrocellulose redox permanganometry is a rapid and simple method for determination of reductive capacity by quantification of the $MnO_2$ precipitate on the nitrocellulose membrane after reducing $KMnO_4$ by a nitrocellulose-prefixed sample. After NRP protocol, the membrane is left to dry. Once sufficiently dry, the NRP membrane is digitalized. Although we suggest that the membrane is digitalized in the first 24 hours, here, we wanted to examine whether the $MnO_2$ precipitate is stable and analyzable even after a prolonged delay. Two NRP membranes were digitalized and analyzed two times with a time delay of 9 months. The same HistoNRP membrane with 4 rat brainstem regions analyzed at baseline and after a 9 month delay is presented in **Fig S4-1A**. Although the NRP signal is still conserved even after 9 months, membrane analysis reveals significant fading and signal equalization. An example of the linear sampling pixel intensity profile plot of the approximately same region is visualized in **Fig S4-1B** with the potential source of problematic ratiometric quantification evident for pixels number 15 and 10 in the 9 month delay analysis with ratiometric values for the corresponding baseline pixels being 3.2 and 2.6 respectively. A membrane with six hippocampal homogenates analyzed with standard NRP is presented in **Fig S4-1C** showing both the baseline signal and the same membrane analyzed after a 9 month period. A significant fading effect was observed. However, in contrast to HistoNRP, time delay in standard NRP resulted in more pronounced differences in contrast to equalization effect. A possible explanation is fading-induced subtraction of the background signal evident on the baseline membrane (**Fig S4-1C**). Integrated density-based quantification of the membrane is presented in **Fig S4-1D** with corresponding ratiometric values (baseline integrated density over time delay integrated density) for individual samples depicted as data point labels next to baseline values. In conclusion, even though meaningful NRP data can be obtained after an enormous time delay (of at least 9 months), we strongly suggest the analysis to be done right after the NRP protocol (in the first 24 hours after the analysis). Time-delay-induced background correction evident in **Fig S4-1C** should be corrected by a longer washing period, shorter incubation in $KMnO_4$, or using standard rolling ball background subtraction in Fiji (NIH, USA) [Process>Subtract Background].

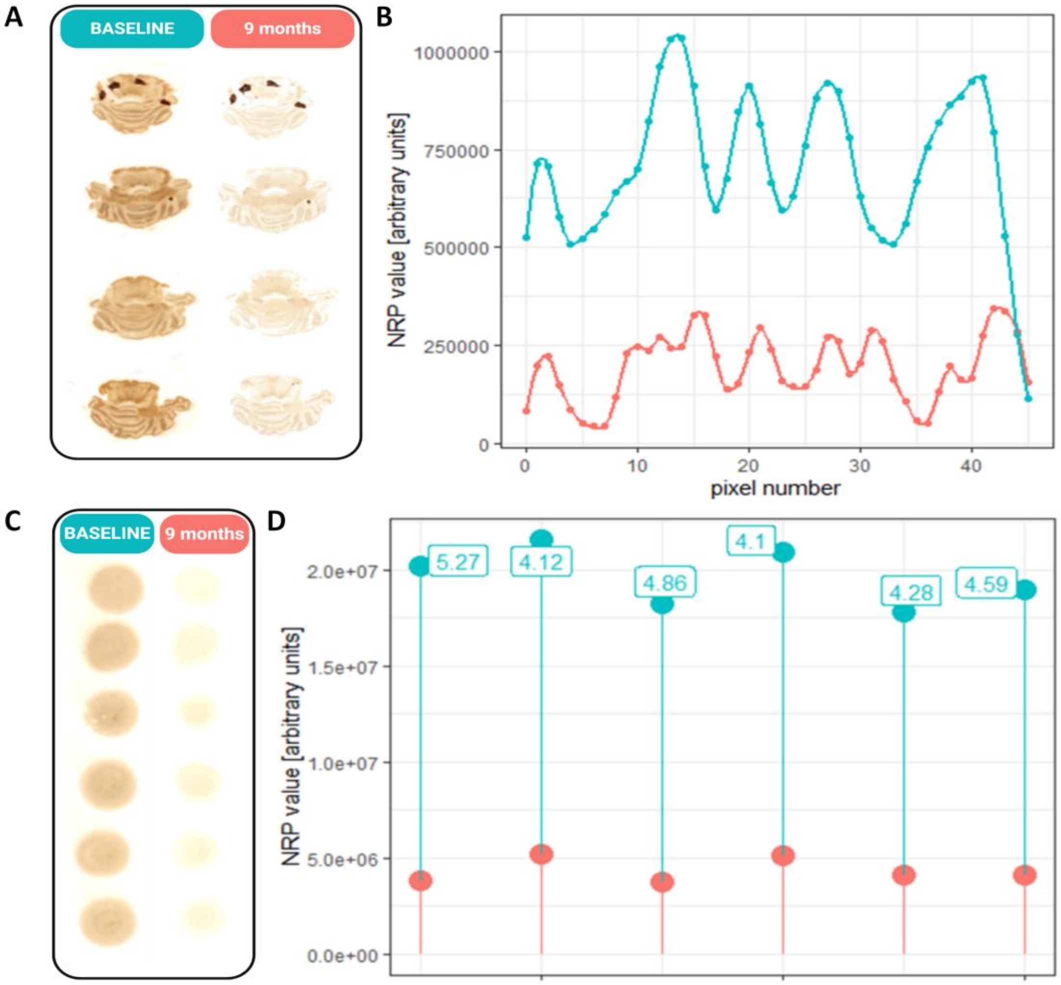


**Fig S4-1. Nitrocellulose redox permanganometry (NRP) membrane stability analysis. A)** HistoNRP membrane with 4 rat brainstem and cerebellum samples digitalized right after the NRP procedure (left) and after a 9 month delay (right). **B)** An example of linear sampling pixel intensity profile plot of the approximately same region right after the NRP procedure (green) and after a 9 month delay (red). **C)** A membrane with six hippocampal homogenates analyzed with a standard NRP right after the NRP procedure (left) and after a 9 month delay (right). **D)** Integrated density-based quantification of the membrane from Fig 1C with baseline values presented in green and values after a 9 month time delay presented in red. Corresponding ratiometric values (baseline integrated density over time delay integrated density) for individual samples are depicted as data point labels next to baseline values.
