## Supplement 5 - Comparison of Nitrocellulose redox permanganometry (NRP) digitalization techniques for "Nitrocellulose redox permanganometry: a simple method for reductive capacity assessment"

Digitalization of the NRP membrane is a prerequisite for reductive capacity assessment based on quantification of the $MnO_2$ precipitate. In the NRP protocol the precipitate is quantified digitally using image analysis; however, the membrane first has to be transferred into a digital form. As with other digitalization techniques (eg. western blot membranes), it is important that the digital image reflects the true objective state of the membrane. For this reason, we analyzed linearity of different digitalization techniques knowing all methods will inevitably suffer from some inherent limitations. For example, a simple office scanner was used for most of the experiments, having in mind that most desktop scanners have both a light source and a sensor on the same side of the scanned object, such as paper or in this case, the nitrocellulose membrane, and a reflecting surface on the other. For this reason, the classic principles of the Beer-Lambert law cannot be assumed as the light from the source will pass twice through the membrane and the photoresistor light entrance angle will vary. Nevertheless, we wanted to make NRP as user friendly as possible, and as precise as possible so various simple digitalization techniques were tested and the best ones were further studied and validated. Four different simple and widely available digitalization techniques are presented in **Fig S5-1.**, and linear regression of the obtained values for a single biological sample diluted in different concentrations. The biological sample used in the experiment was a single hippocampal homogenate of a control animal with protein concentration of 15.61 µg/µl, as determined by the Lowry assay [1]. Raw images of NRP membranes digitalized with different instruments and corresponding density plots obtained by the Gel Analyzer plugin are shown in **Fig S5-1A-D.** An NRP membrane digitalized by RICOH SP 4410SF desktop office scanner is shown in **Fig S5-1A**, the same membrane digitalized by a Samsung Galaxy S8 cell phone camera with a flash option turned on is presented in **Fig S5-1B,** or with a flash option turned off in **Fig S5-1C**. Additionally, the membrane was digitalized with a MicroChemi (DNR Bio Imaging Systems, Israel) the camera at -56°C, standardly used for visualization of chemiluminescence with illumination turned on (**Fig S5-1D**). Linear regression with corresponding equations, Pearson correlation coefficients and p values is shown in **Fig S5-1E**. Based on validation experiments for different digitalization techniques, the office scanner digitalization was chosen as the best method, although Beer-Lambert principles cannot be directly assumed, the method is inherently defined by the enclosed digitalization protocol, important for maximal reduction of error induced by environmental illumination variability, results are rapidly available, and the method shows good linearity. Results obtained with a Samsung Galaxy S8 cell phone were also very reliable, and even slightly more sensitive in comparison to the scanner. However, considering potential illumination variability differences, we decided to standardly rely on the office scanner for digitalization of our NRP membranes.

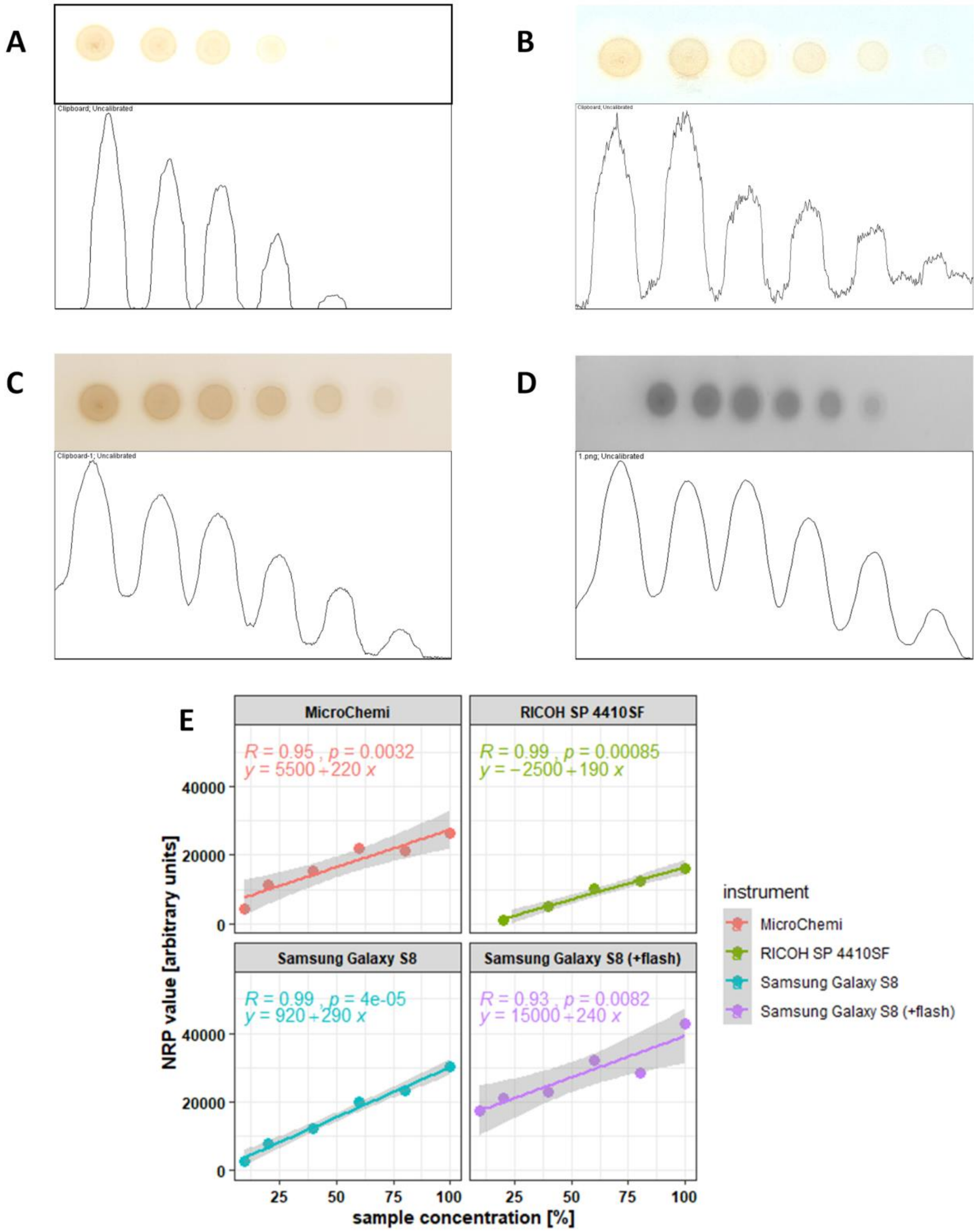


**Fig S5-1.** Linearity comparison for digitalization with different instruments. A single NRP membrane obtained by analysis of different dilutions of a single hippocampal homogenate from the control rat (1,0.8, 0.6, 0.4, 0.2, 0.1). **A)** An NRP membrane and corresponding density plot digitalized by the RICOH SP 4410SF desktop office scanner. **B)** An NRP membrane and corresponding density plot digitalized by a Samsung Galaxy S8 cell phone camera with a flash option turned on. **C)** An NRP membrane and corresponding density plot digitalized by a Samsung Galaxy S8 cell phone camera with the flash option turned off. **D)** An NRP membrane and corresponding density plot digitalized by the MicroChemi (DNR Bio Imaging Systems, Israel) camera at -56°C, standardly used for visualization of chemiluminescence with illumination turned on. **E)** Linear regressions with corresponding equations, Pearson correlation coefficients, and p values for data obtained following digitalization with different instruments. All density plots were obtained by the Gel Analyzer plugin for Fiji (NIH, USA) and data was extracted by measuring the area under the curve.
