## Supplement 6 - An explanation of the computational analysis of Nitrocellulose redox permanganometry (NRP) for "Nitrocellulose redox permanganometry: a simple method for reductive capacity assessment"

To obtain accurate results of sample reductive capacity, adequate digitalization and analysis of the membrane is indispensable. This document provides a step-by-step protocol on how to analyze NRP membranes to ensure proper implementation of NRP into research practice. A single NRP membrane digitalized by RICOH SP 4410SF desktop office scanner as described in **Supplement 6** was used here for demonstrative purposes. Although different software solutions can be used for signal quantification, we recommend Fiji (Fiji Is Just ImageJ), a version of the popular ImageJ software (NIH, USA) with pre-installed plugins. Precise quantification of the NRP signal can be simply done by using the Fiji Gel Analyzer plugin and following the same protocol as recommended for the dot blot analysis. In the following protocol the steps are shown in green and the corresponding ImageJ macro code is shown in red.

---

1) Import the membrane image into Fiji.
   [File>Open…]
   [open("D:/Users/jan.homolak/Desktop/SampleNRP.png");]

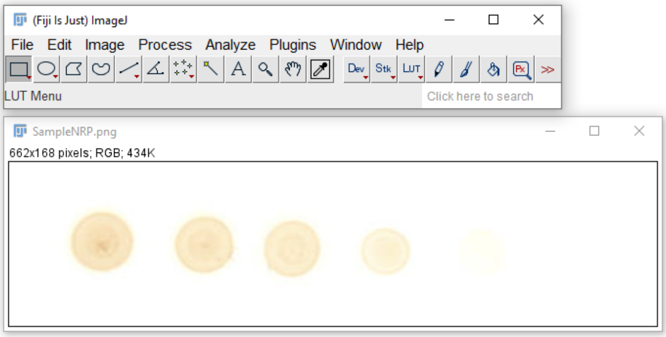


2) Use the rectangle tool to select samples
   [Rectangle]
   [//setTool("rectangle");
   makeRectangle(9, 19, 602, 131);]

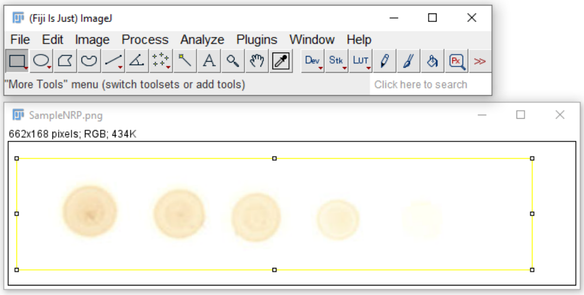


3) Use the Gel Analyzer "Select First Lane" option
   [Analyze>Gels>Select First Lane//OR//CTRL+1]

[run("Select First Lane");]

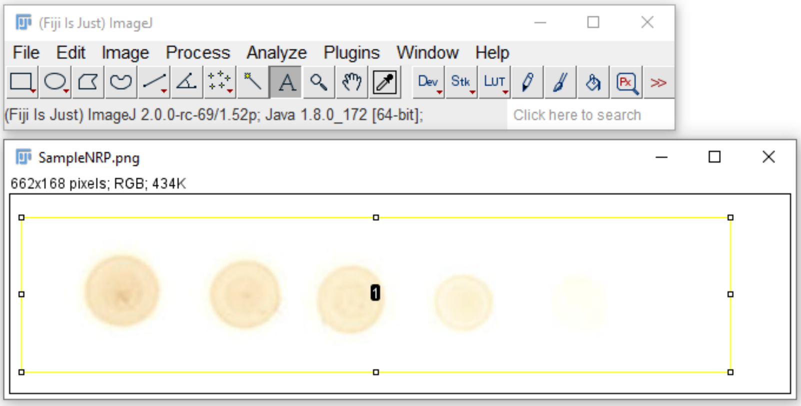


4) Use the Gel Analyzer "Plot Lanes" option [Analyze>Gels>Plot Lanes//OR//CTRL+3] [//setTool("line"); run("Plot Lanes");]

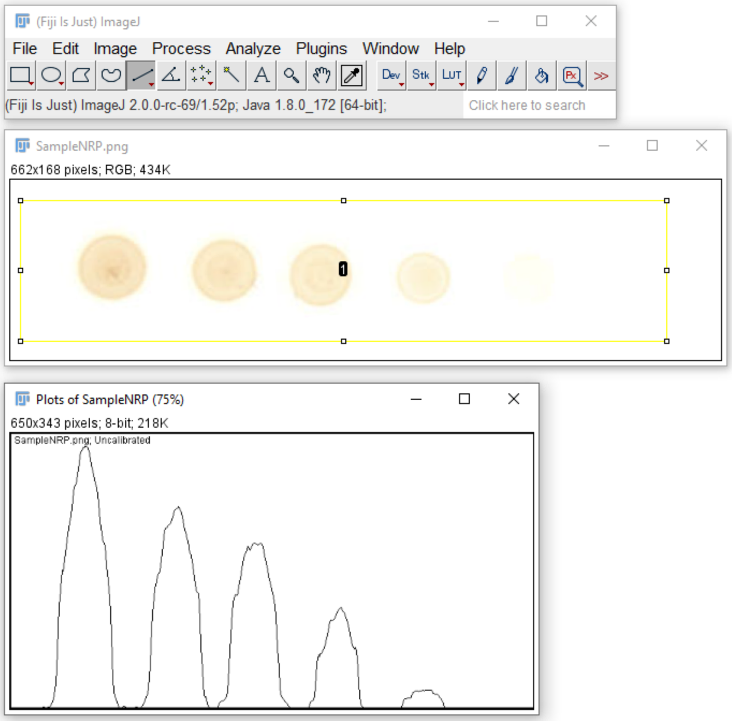


5) Use the Wand tool to select density plots of the corresponding samples and calculate area under the curve [Wand tool]

[//setTool("wand");
doWand(94, 189);
doWand(209, 253);
doWand(298, 220);
doWand(420, 282);
doWand(518, 328);]

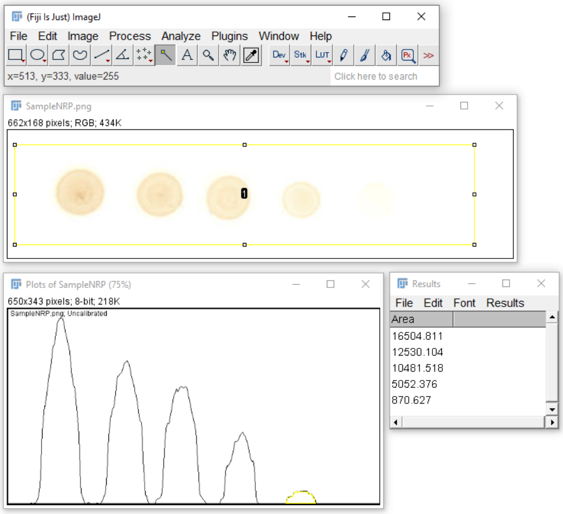


6) Select the values in the Results window and save the results for subsequent analysis and visualization. Alternatively, you can select the results with the pointer and copy and paste them into an excel sheet [Results:File>Save As...//OR//CTRL+S] [saveAs("Results", "D:/Users/jan.homolak/Desktop/SampleNRPresults.csv");]

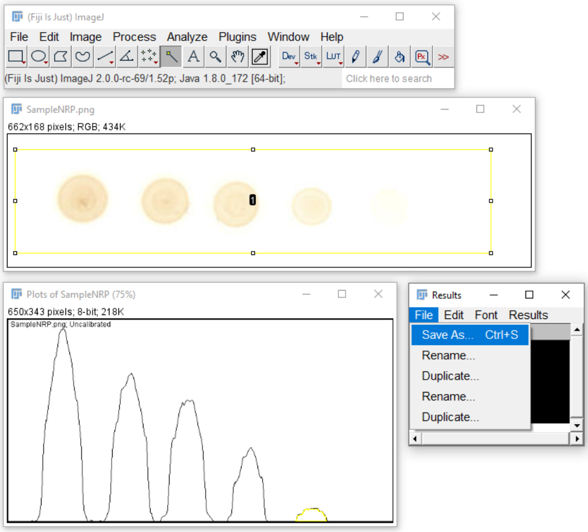


7) Alternatively, you can directly plot the results inside Fiji. [Results:Results>Plot…] [Plot.create("Plot of Results", "x", "Area"); Plot.add("Connected Circles", Table.getColumn("Area", "Results"));

Plot.setStyle(0, "red,#a0a0ff,2.0,Connected Circles");]

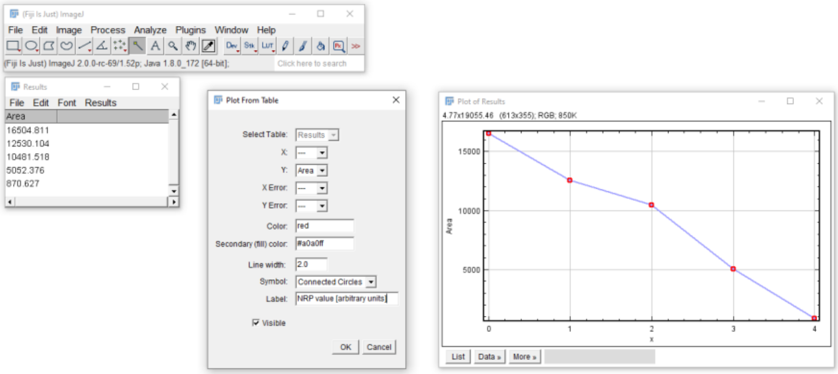


8) Further analysis can also be done directly on the obtained results. An example of regression analysis is shown.
[Plot of Results:Data>>Add fit]
[Plot.setStyle(2, "red,#a0a0ff,2.0,Line");]

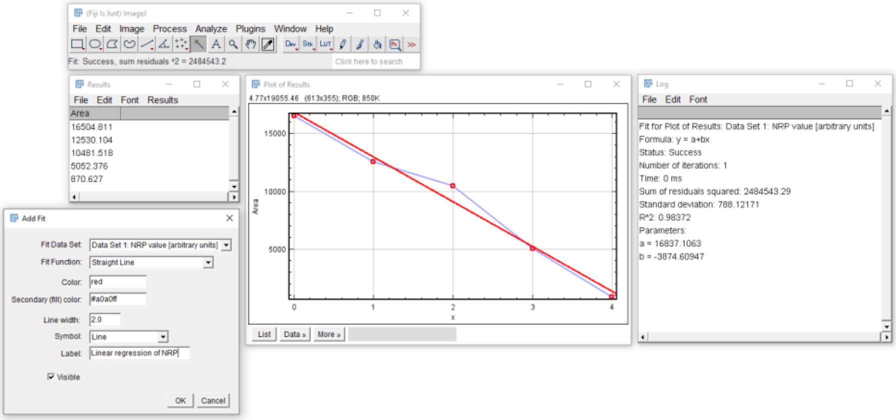


9) Additional visual 3D representation of the NRP signal can also be done in Fiji by using the 3D Surface Plot Analysis option
[File>Open…
Analyze>3D Surface Plot]
[run("3D Surface Plot");]

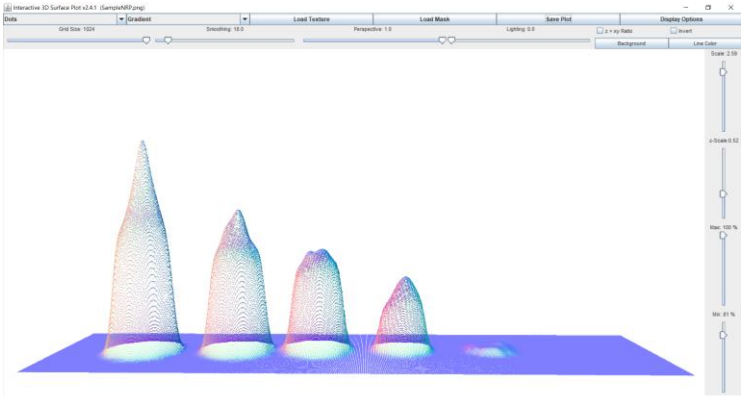
