## Supplement 7 - Additional replications of ORP-NRP experiments from Fig 1E and Fig 1F for "Nitrocellulose redox permanganometry: a simple method for reductive capacity assessment"

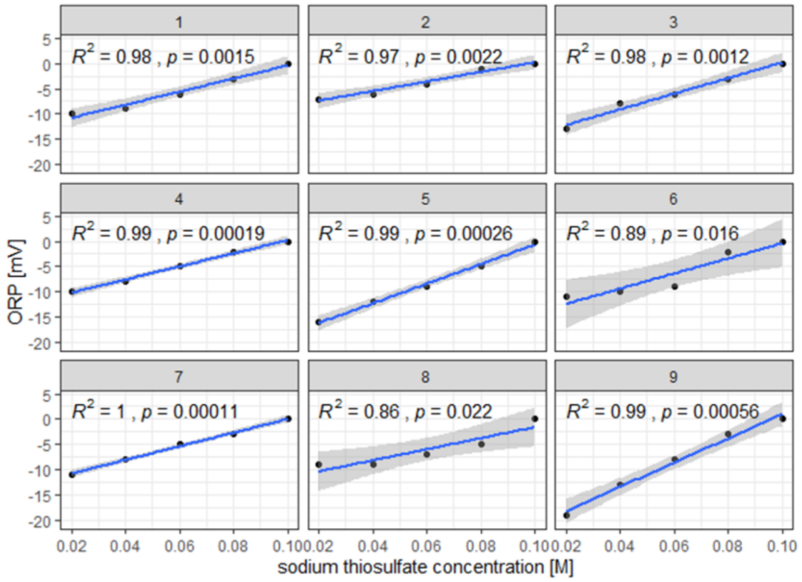


**Fig S7-1.** Additional replications of the experiment presented in Fig 1E in the Main text.

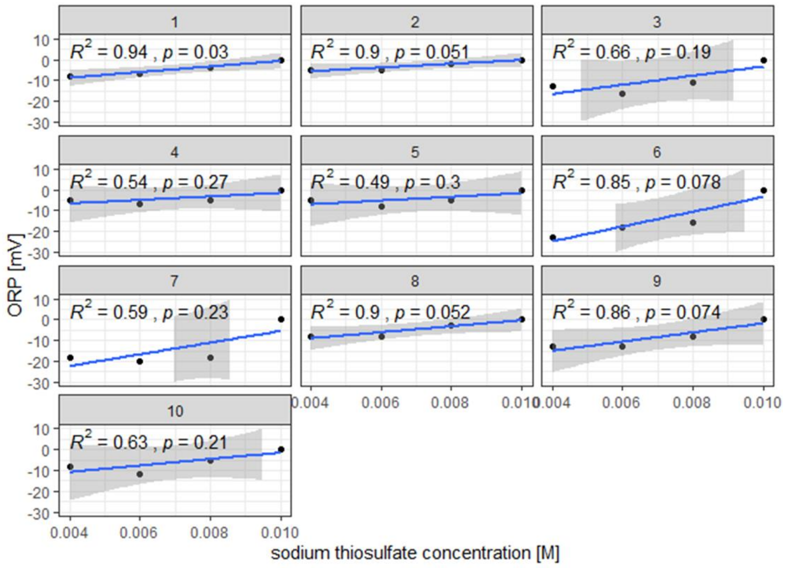


**Fig S7-2.** Additional replications of the experiment presented in Fig 1F in the Main text.
