## Supplement 8 - A detailed explanation of the origin of animal tissue used in the Nitrocellulose redox permanganometry (NRP) proof of concept experimen for "Nitrocellulose redox permanganometry: a simple method for reductive capacity assessment"

### A detailed explanation of the origin of animal tissue used in the Nitrocellulose redox permanganometry (NRP) proof of concept experiments

In short, animal tissue from 3 different animal experiments was used in the NRP proof of concept studies. A single hippocampal homogenate from a single control animal was used for the analysis of exogenous redox system manipulation of the biological specimen illustrated in the main text in the **Fig 1F**. The sample we used was obtained from **Experiment 1**. The sample was used in **Supplement 6** (A comparison of NRP membrane digitalization techniques) and in **Supplement 10** (Protein content correction analysis). Plasma and hippocampal homogenates obtained from 6 control rats and 6 rats treated intracerebroventricularly with streptozotocin (STZ-icv) was used for proof of concept of biological specimen NRP analysis presented in **Fig 2A** and **Fig 2C**. Plasma and hippocampal homogenates from 6 wild type mice and 6 transgenic tg2576 animals was used for additional proof of concept of NRP analysis of biological samples. These samples were obtained in **Experiment 2**. Finally, tissue used for demonstration of HistoNRP shown in **Fig 2F-J**, as well as tissue used for NRP membrane stability analysis shown in **Supplement 4** was harvested in the **Experiment 3**. Here, 6 hippocampal homogenates of untreated rats were used (**Supplement 4**), as well as tissue sections of the brain stem region from 4 animals (**Supplement 4**) and 1 coronal section of the rat brain damaged unilaterally with a microdialysis probe (**Main text Fig 2F-J**).

#### Experiment 1:

The experiment aimed to assess acute effects of oral and intraperitoneal galactose load in a rat model of sporadic Alzheimer's disease. Forty male Wistar rats, bred and kept in-house (Department of Pharmacology, University of Zagreb School of Medicine) were used for the experiment. The animals were kept in an animal facility with stable conditions (constant temperature and humidity – 22-24ºC and 40-60%, and 12-h light/12-h dark cycle) in standard cages with wood-chip bedding and food and water ad libitum; with three animals placed per cage. At the age of three months, the animals underwent intracerebroventricular treatment with either streptozotocin (STZ-icv; 30 animals) or 0.05M citrate buffer, pH 4.5 (CTR; 10 animals). STZ is a beta-cytotoxic substance selective for insulin-secreting/producing cells, which has been abundantly used to induce insulin-resistant brain state in laboratory animals, mimicking the main hallmarks of sporadic Alzheimer's disease. STZ was applied intracerebroventricularly to rats in deep anesthesia (ketamine 50 mg/kg / xylazine 5 mg/kg, ip) in a dose of 3 mg/kg divided in two doses (48 hours apart) into the rats' lateral ventricles (according to the procedure first described by Noble et al. [1] and used by our research team in previous experiments [2–4], whereas control animals received only the vehicle, citric buffer, in the same manner. One month after STZ-icv treatment, 20 STZ-icv treated rats were divided into 2 groups, with 10 animals per group, and received either a single oral dose of galactose via gastric tube (200 mg/kg dissolved in 1 mL of water, equivalent to a 6% galactose solution) or a single ip dose (200 mg/kg dissolved in 1 mL of saline solution, equivalent to a 6% galactose saline solution), adjusted according to the rats' body weight. The remaining 10 CTR and 10 STZ animals were left as intact controls. Fifteen minutes after the galactose load (time point chosen based on literature data [5] due to rapid conversion of galactose to glucose in blood), the animals were anesthetized with a combination of ketamine and xylazine and blood samples were taken from the retro-orbital sinus. Whole blood samples were collected to heparinized tubes, centrifuged at 3600 RPM for 10 minutes and the resulting supernatants frozen and stored at -80ºC. Rats were then sacrificed in two ways, pertaining to the purpose of the collected tissue. Four animals per group, while still in deep anesthesia, underwent transcardial fixative perfusion [6], where the tissue is first washed with saline and subsequently fixated with 4% buffered paraformaldehyde. Upon removal of the intestines and brains, the tissue is submerged in a 15% sucrose-buffered paraformaldehyde solution followed by 30% sucrose-phosphate buffer solution immersion and consequent freezing at -80ºC for long-term storage. The remaining six animals per group underwent cervical dislocation and their intestines and brains were quickly removed and dissected on ice, with removal of the hippocampi, freezing using liquid nitrogen and storage at -80ºC. Hippocampal samples were later thawed and ultrasonically homogenized in a lysis buffer containing 50 mM Trizma base (pH 8), 150 mM sodium chloride, 0,5 mM EDTA, 1 mM dithiothreitol, 0,01 M sodium vanadate, 0,5% sodium deoxycholate, 1% NP-40 detergent and 0,1% SDS. The sonicated samples were then centrifuged at 12000 RPM for 10 minutes, and total protein content from the separated supernatants was measured using the Lowry method [7]. Finally, supernatants were stored at -80ºC until further analysis.

#### Experiment 2:

The experiment was devised to examine the potential therapeutic effects of oral galactose treatment in a mouse model of familial Alzheimer's disease. Twenty adult male B6; SJL-Tg(APPSWE)2576 Kha heterozygous transgenic mice (TG) overexpressing the human amyloid precursor protein (APP), and 20 corresponding wild types (WT) were purchased from Taconic Biosciences Inc. (Hudson NY, USA). From the age of 3 months onwards, mice were individually housed in appropriate cages with wood-chip bedding, which were located in ventilated cabinets to ensure stable temperature (22–24 °C) and humidity (40–60%) conditions. The cabinets were placed in an air-conditioned room with a 12-h light/12-h dark cycle, specialized for housing of transgenic animals (Croatian Institute for Brain Research, University of Zagreb School of Medicine). At the age of 10 months, when, according to literature data [8,9], this model develops cognitive deficits and neurochemical changes, but no plaque accumulation is evident, the animals entered the experiment, and baseline cognitive and metabolic parameters were measured before initiation of galactose therapy. Cognitive function was assessed using the Morris Water Maze (MWM) swimming test for spatial learning [10], nesting behavior was evaluated by scoring the quality of the animals' nest formation from the given material per standard protocol [11], glucose metabolism and homeostasis were measured by intraperitoneal glucose tolerance testing (ipGTT) in fasting mice [12] and anxiety-related behavior was assessed by standard elevated plus maze test (EPM) [13]. After the establishment of baseline values, mice were divided equally into 4 groups based on cognitive results, with 10 animals per group. One TG and one WT group started 2-month oral galactose therapy, whereas the remainder of animals kept receiving tap water ad libitum. Oral galactose (Sigma Aldrich, USA) was dissolved in tap water at a dose of 200 mg/kg/day; a dose selected based on previous experiments where it successfully prevented/normalized early cognitive decline in an intracerebroventricular-streptozotocin model of sporadic Alzheimer's disease in rats [14,15]. After 2 months of oral galactose treatment, mice underwent MWM, passive avoidance testing (PAT) for aversive memory assessment [16], nesting-behavior assessment, ipGTT, EPM, open field testing (OF) for locomotor and anxiety-like behavior [17] and in vivo brain glucose uptake measurement of 18fluorodeoxyglucose by PET scan (FDG-PET; 4 animals per group). Upon the completion of cognitive and metabolic tests, mice were anesthetized with a combination of ketamine and xylazine and blood samples were taken from the retro-orbital sinus. Whole blood samples were collected to heparinized tubes, centrifuged at 3600 RPM for 10 minutes and the resulting supernatants frozen and stored at -80ºC. Mice were then sacrificed in two ways, pertaining to the purpose of the collected tissue. Four animals per group, while still in deep anesthesia, underwent transcardial fixative perfusion [6], where the tissue is first washed with saline and subsequently fixated with 4% buffered paraformaldehyde. Upon removal of the intestines and brains, the tissue is submerged in a 15% sucrose-buffered paraformaldehyde solution followed by 30% sucrose-phosphate buffer solution immersion and consequent freezing at -80ºC for long-term storage. The remaining six animals per group underwent cervical dislocation and their intestines and brains were quickly removed and dissected on ice, with removal of the hippocampi, freezing using liquid nitrogen and storage at -80ºC. Hippocampal samples were later thawed and ultrasonically homogenized in a lysis buffer containing 50 mM Trizma base (pH 8), 150 mM sodium chloride, 0,5 mM EDTA, 1 mM dithiothreitol, 0,01 M sodium vanadate, 0,5% sodium deoxycholate, 1% NP-40 detergent and 0,1% SDS. The sonicated samples were then centrifuged at 12000 RPM for 10 minutes, and total protein content from the separated supernatants was measured using the Lowry method [7]. Finally, supernatants were stored at -80ºC until further analysis.

**Experiment 3:**

The experiment aimed to assess acute metabolic and redox regulatory network time-response (up to 2h) after oral galactose load in rats. Forty male Wistar rats, bred and kept in-house (Department of Pharmacology, University of Zagreb School of Medicine) were used for the experiment. The animals were kept in an animal facility with stable conditions (constant temperature and humidity – 22-24ºC and 40-60%, and 12-h light/12-h dark cycle) in standard cages with wood-chip bedding and food and water ad libitum; with three animals placed per cage. At the age of 3 months, the animals were randomized into 5 groups as follows: baseline control group (CTR; n=8), 3 galactose treatment groups sacrificed either in 0.5h time point (n=8), 1h time point (n=8), 2h time point (n=8), and an additional group also sacrificed 2 hours after the galactose treatment but with a preinstalled microdialysis probe (n=8). The microdialysis probe was installed 24 hours before the experiment. In short, rats were anesthetized with isoflurane in the Isoflurane Vaporizer (Ugobasile, Italy) and placed in the stereotaxic apparatus (Stoelting, USA). A microdialysis probe (CMA Microdialysis AB, Sweden) was placed and fixed with GC FujiCEM (GC Corporation, Japan). 50 mg/ml of metamizole was administered for postsurgical analgesia. On the day of the experiment, rats either received no treatment (CTR) or received 200 mg/kg of galactose dissolved in 1 mL of water by orogastric gavage and were sacrificed either 0.5, 1 or 2 hours after the treatment. Animals were anesthetized with a combination of ketamine and xylazine and blood samples were taken from the retro-orbital sinus. Whole blood samples were collected to heparinized tubes, centrifuged at 3600 RPM for 10 minutes and the resulting supernatants frozen and stored at -80ºC. Rats were then sacrificed in two ways, pertaining to the purpose of the collected tissue. Four animals per group, while still in deep anesthesia, underwent transcardial fixative perfusion [6], where the tissue is first washed with saline and subsequently fixated with 4% buffered paraformaldehyde. Upon removal of the intestines and brains, the tissue is submerged in a 15% sucrose-buffered paraformaldehyde solution followed by 30% sucrose-phosphate buffer solution immersion and consequent freezing at -80ºC for long-term storage. The remaining six animals per group underwent cervical dislocation and their intestines and brains were quickly removed and dissected on ice, with removal of the hippocampi, freezing using liquid nitrogen and storage at -80ºC. Hippocampal samples were later thawed and ultrasonically homogenized in a lysis buffer containing 50 mM Trizma base (pH 8), 150 mM sodium chloride, 0,5 mM EDTA, 1 mM dithiothreitol, 0,01 M sodium vanadate, 0,5% sodium deoxycholate, 1% NP-40 detergent and 0,1% SDS. The sonicated samples were then centrifuged at 12000 RPM for 10 minutes, and total protein content from the separated supernatants was measured using the Lowry method [7]. Finally, supernatants were stored at -80ºC until further analysis.

Some of the results from Experiment 1 and Experiment 2 were published in [14,15].

1. Noble, E. P., Wurtman, R. J. & Axelrod, J. A simple and rapid method for injecting H3-norepinephrine into the lateral ventricle of the rat brain. *Life Sci.* **6**, 281–291 (1967).
2. Knezovic, A. *et al.* Staging of cognitive deficits and neuropathological and ultrastructural changes in streptozotocin-induced rat model of Alzheimer's disease. *J. Neural Transm.* **122**, 577–592 (2015).
3. Grünblatt, E., Salkovic-Petrisic, M., Osmanovic, J., Riederer, P. & Hoyer, S. Brain insulin system dysfunction in streptozotocin intracerebroventricularly treated rats generates hyperphosphorylated tau protein. *J. Neurochem.* **101**, 757–770 (2007).
4. Salkovic-Petrisic, M. *et al.* Cerebral amyloid angiopathy in streptozotocin rat model of sporadic Alzheimer's disease: a long-term follow up study. *J. Neural Transm.* **118**, 765–772 (2011).
5. Kliegman, R. M. & Morton, S. Sequential intrahepatic metabolic effects of enteric galactose alimentation in newborn rats. *Pediatr. Res.* **24**, 302–307 (1988).
6. Gage, G. J., Kipke, D. R. & Shain, W. Whole animal perfusion fixation for rodents. *J. Vis. Exp.* (2012) doi:10.3791/3564.
7. Lowry, O. H., Rosebrough, N. J., Farr, A. L. & Randall, R. J. Protein measurement with the Folin phenol reagent. *J. Biol. Chem.* **193**, 265–275 (1951).
8. Oddo, S., Caccamo, A., Smith, I. F., Green, K. N. & LaFerla, F. M. A dynamic relationship between intracellular and extracellular pools of Abeta. *Am. J. Pathol.* **168**,

184–194 (2006).

9. Lesné, S. *et al.* A specific amyloid-beta protein assembly in the brain impairs memory. *Nature* **440**, 352–357 (2006).
10. Vorhees, C. V. & Williams, M. T. Morris water maze: procedures for assessing spatial and related forms of learning and memory. *Nat. Protoc.* **1**, 848–858 (2006).
11. Deacon, R. M. J. Assessing nest building in mice. *Nat. Protoc.* **1**, 1117–1119 (2006).
12. Ayala, J. E. *et al.* Standard operating procedures for describing and performing metabolic tests of glucose homeostasis in mice. *Dis. Model. Mech.* **3**, 525–534 (2010).
13. Walf, A. A. & Frye, C. A. The use of the elevated plus maze as an assay of anxiety-related behavior in rodents. *Nat. Protoc.* **2**, 322–328 (2007).
14. Knezovic, A. *et al.* Glucagon-like peptide-1 mediates effects of oral galactose in streptozotocin-induced rat model of sporadic Alzheimer's disease. *Neuropharmacology* **135**, 48–62 (2018).
15. Babic Perhoc, A. *et al.* Cognitive, behavioral and metabolic effects of oral galactose treatment in the transgenic Tg2576 mice. *Neuropharmacology* **148**, 50–67 (2019).
16. Walters, G. C. & Abel, E. L. Passive avoidance learning in rats, mice, gerbils, and hamsters. *Psychon. Sci.* **22**, 269–270 (1971).
17. Seibenhener, M. L. & Wooten, M. C. Use of the Open Field Maze to measure locomotor and anxiety-like behavior in mice. *J. Vis. Exp.* e52434 (2015).
