## Supplement 9 - Nitrocellulose redox permanganometry (NRP) protein concentration-based correction analysis for "Nitrocellulose redox permanganometry: a simple method for reductive capacity assessment"

Biological sample concentration correction is often done by normalization to total protein content. A simple ratiometric correction assumes a linear relationship between measured properties and protein content. However, this assumption is rarely tested directly. In order to make sure ratiometric correction for protein content is a valid approach for normalization of NRP values obtained from biological samples with different protein concentrations we designed an experiment to test linearity between protein concentration and NRP values. In short, a single biological sample (hippocampal homogenate from a single control rat from **Experiment 1**) with protein concentration of 15.61 µg/µl as determined by folin phenol protein quantification [1]was diluted at 80, 60, 40, 20 and 10% of its original concentration with dd$H_2O$ to make a set of samples for linearity comparisons. Samples (1 µl) were pipetted onto the nitrocellulose membrane in duplicates and the membranes were left to dry. Once dry, one membrane was stained with the 0.1% (w/v) Ponceau S in 5% (v/v) acetic acid [2,3] for additional protein quantification, and standard NRP protocol was used on the other membrane. Both membranes were scanned and analyzed in Fiji with the Gel Analyzer plugin (NIH, USA). Analysis of measured and predicted NRP values following the principle of maximal linearity and corresponding percent deviations are illustrated in **Fig S9-1A.** As evident from the graph even from the protein concentrations equal to 40% of the original concentration, deviation from maximal linearity was just 12.7% so we concluded a ratiometric approach is justified for protein concentrations deviating from the standard concentration by approximately 50% of the nominal value. Next we were interested in the source of variation to see whether dispersion increasing with a factor of deviation from the nominal point is inherently related to NRP, or to the principle of nitrocellulose digital quantification analysis so we compared the values obtained with NRP to the values of signal intensity quantification for the same samples analyzed with Ponceau S nitrocellulose protein staining (**Fig S9-1B**). Interestingly, NRP values demonstrated better linearity and less dispersion from the absolute linear principles in comparison with the Ponceau S nitrocellulose protein quantification.

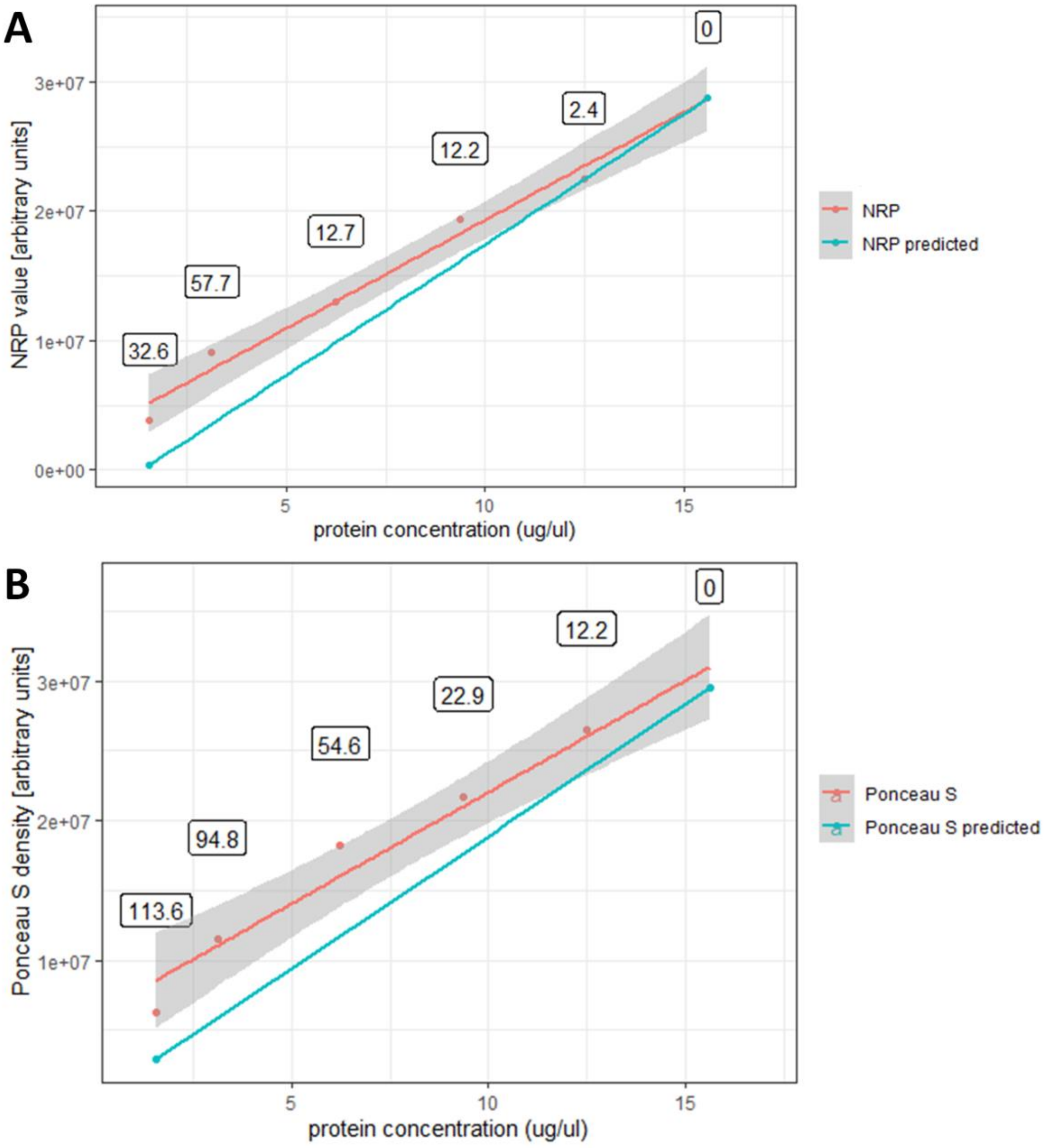


**Fig S9-1.** Comparison of predicted and obtained NRP (**A**) and Ponceau S (**B**) values from serial dilutions of a biological sample.

1. Lowry, O. H., Rosebrough, N. J., Farr, A. L. & Randall, R. J. Protein measurement with the Folin phenol reagent. *J. Biol. Chem.* **193**, 265–275 (1951).

2. Bannur, S. V., Kulgod, S. V., Metkar, S. S., Mahajan, S. K. & Sainis, J. K.

Protein determination by ponceau S using digital color image analysis of protein spots on nitrocellulose membranes. *Anal. Biochem.* **267**, 382–389 (1999).

3. Sander, H., Wallace, S., Plouse, R., Tiwari, S. & Gomes, A. V. Ponceau S waste: Ponceau S staining for total protein normalization. *Anal. Biochem.* **575**, 44–53 (2019).
