## Supplement 10 - HistoNRP adaptation for the analysis of formalin-fixed paraffin-embedded (FFPE) tissue sections for "Nitrocellulose redox permanganometry: a simple method for reductive capacity assessment"

Spatial reductive capacity of tissue sections by HistoNRP method depends on the success of molecular content of the tissue onto the nitrocellulose membrane. In order to ensure optimal transfer from the FFPE sections, a standard HistoNRP protocol explained in the Supplement 1 has to be adapted. The main difference between HistoNRP for cryosections and FFPE HistoNRP is an additional trypsin-based enzymatic retrieval step, and introduction of heat-facilitated passive diffusion slice printing step instead of classic passive diffusion slice print blotting used for the transfer of cryosections onto the nitrocellulose membrane. In short, FFPE tissue is first cut on the microtome and tissue sections are mounted onto the histological slides and processed by the standard deparaffinization protocol through Xylene followed by decreasing concentrations of ethanol (EtOH)(3x5 min Xylene/ 2x5 min 100% EtOH/ 2x5 min 95% EtOH/ 2x5 min 70% EtOH/ 2x5 min 50% EtOH). Finally, sections are immersed in phosphate buffered saline (PBS; pH 7.4) 2 times for 5 minutes. After PBS, the sections are placed in warm (37°C) antigen retrieval solution (0.05% Trypsin, 0.1% CaCl in ddH$_2$O; pH 7.8) for 45 minutes. After the retrieval, the sections are placed in PBS for 2x5 min at room temperature (RT) to equilibrate. After equilibration, a glass plate is placed on the laboratory heater and the sections are placed on top of the glass plate and wetted with PBS. A piece of nitrocellulose membrane is placed on top of the tissue sections, and 3 filter papers are placed on the nitrocellulose membrane and wetted with PBS. Finally, a piece of parafilm is placed on top of wet filter papers and the edges are pressed towards the bottom glass plate. Additional glass plate is placed on top of the parafilm, and weight (beaker filled with water) is placed on the upper plate to ensure the pressure of approximately 31.384 mmHg (**Fig S10-1**). The heater is set at 60°C, and the timer at 8 hours. Once heating is started the parafilm should be pressed down towards the lower glass plate to ensure optimal isolation and humidity during the transfer protocol. After 8 hours the heater is turned off and the beaker and the upper glass plate are removed carefully. Parafilm and 3 filter papers are removed with tweezers, and nitrocellulose is wetted with PBS, removed, and left to dry. Once dry the membrane can be analyzed with the standard NRP procedure. In order to enable NRP analysis of FFPE, several protocols were designed and tested and the protocol described above gave the best results both in the context of protein transfer (**Fig S10-2A**) and NRP results (**Fig S10-2B**). An image of NRP analysis of the membrane obtained with one protocol without enzymatic retrieval step and heating-facilitated transfer is shown in **Fig S10-2C** for comparison. Nevertheless, several questions remain to be answered and are in focus of our ongoing research. First, we hypothesize that the heating protocol can oxidize the tissue samples, and therefore introduce error in reductive capacity analysis of FFPE tissues by the NRP (or any other method focused on the assessment of FFPE tissue reductive capacity). Robust and detailed experiments have to be designed in order to provide the answer to this question, and help us understand the significance of heat-induced tissue oxidation in this context. Moreover, In the original protocol, we tested different enzymatic retrieval protocols, and different buffers, and good results were obtained by following deparaffinization and enzymatic retrieval protocol for proteomic expressional profiling of FFPE tissue by multiplex immunoblotting proposed by Chung and Hewitt [1]. Their protocol proposes the equilibration step in 50 mM ammonium bicarbonate buffer (pH 8.2) followed by treatment with freshly prepared enzyme cocktail solution that contains 0.001% trypsin plus proteinase K in the same buffer. In their protocol protease inhibition step is added after the enzymatic retrieval and they propose a 15 minute long treatment with proBuffer (One tablet of complete protease inhibitor 0.5 ml of phosphatase inhibitor I and 0.5 ml of phosphatase inhibitor II in PBS (pH 7.2)) at room temperature. Nevertheless, in our preliminary experiments, enzymatic retrieval with greater concentrations of trypsin provided better results so we decided to follow this protocol. In regards to the protease inhibition step, our preliminary data suggest that results remain the same if this step is omitted so we didn't place a protease inhibition step in our protocol although we initially planned. We hypothesize that proteases could still be at least partially inactivated, and heat could further facilitate this inactivation during the transfer protocol. As in our FFPE modification of HistoNRP, the heat-facilitated passive diffusion slice printing step follows the enzymatic retrieval, we consider this step to be optional based on our analyses. Nevertheless, we wanted to emphasize this as the end user should be aware that protein degradation might take place during the protocol, and this might turn out to be important if the protocol is adapted or modified, and the tissue is not heated and transferred immediately. As briefly mentioned above, possible modifications and further understanding of this modified HistoNRP protocol for FFPE still remains an active area of research in our laboratory, so we believe further experiments might provide new information and result in further modifications of this protocol.

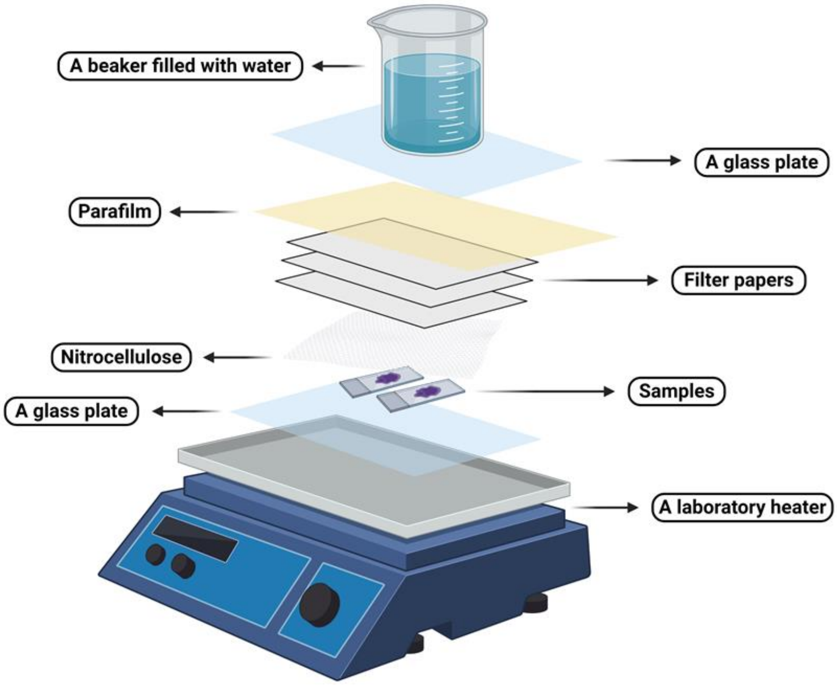


**Fig S10-1.** A schematic representation of the heat-facilitated passive diffusion slice printing setup.

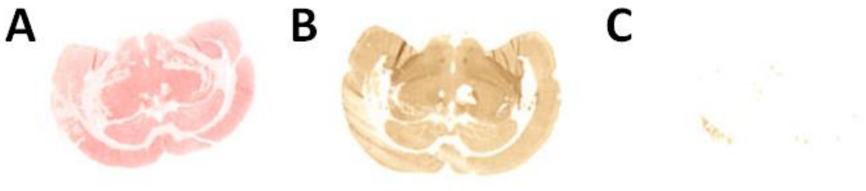


**Fig S10-2.** Analysis of FFPE brain tissue fixed onto the membrane by heat-facilitated passive diffusion slice printing. A) Ponceau S analysis of proteins transferred to the nitrocellulose membrane by heat-facilitated passive diffusion slice printing. B) An example of the FFPE brain tissue analyzed by NRP after heat-facilitated passive diffusion slice printing. C) An example of NRP analysis of the membrane obtained by passive diffusion slice print blotting from the standard NRP protocol instead of trypsin-based retrieval followed by heat-facilitated passive diffusion slice printing.

1. Chung, J.-Y. & Hewitt, S. M. Proteomic expressional profiling of a paraffin-embedded tissue by multiplex tissue immunoblotting. *Methods Mol. Biol.* **1312**, 175–184 (2015).
