## Supplement 11 - A detailed explanation of the HistoNRP demonstration analysis illustrated in the Fig 2 of the Main text for "Nitrocellulose redox permanganometry: a simple method for reductive capacity assessment"

A coronal section of the rat brain damaged unilaterally with the microdialysis probe was used for demonstration of the spatial distribution analysis of tissue reductive capacity illustrated in the Main text in **Fig 2**. A microdialysis experiment is explained in detail in **Supplement 9**. A standard HistoNRP protocol (as described in **Supplement 1**) was used for reduction capacity analysis and the membrane was digitalized both with an office scanner and with Samsung Galaxy S8. An image obtained by the Samsung Galaxy S8 camera was used for detailed analysis as it provided more anatomical details. Rat brain stereotaxic atlas was used to identify anatomical areas of interest [1] and Fiji software was used to calculate distribution of pixel intensities in the inverted 8-bit image. In short, an image was imported into Fiji [File>Open…], converted to 8-bit [Image>Type>8-bit], and inverted [Edit>Invert]. First, the brain was divided by 6 lines of interest as shown in **Fig 2H** in the Main text and intensity profiles were calculated for both ipsilateral and the contralateral side using the straight line selection tool followed by the plot profile option [Analyze>Plot Profile]. Data was exported as a list, saved in an excel sheet and imported in R software environment for statistical computing for further analysis and visualization with the ggplot2 package (**Main text Fig 2I**). Moreover, anatomical areas of interest were defined on the ipsilateral and contralateral side of the brain using the selection tool of the same size, and pixel intensity value distribution was calculated using the Histogram option [Analyze>Histogram ]. Values were exported as a list, and saved in an excel sheet in the format of a list of pixel intensities with corresponding number of pixels for the individual intensity value. Additionally, pixel distance was calculated for all areas of interest. Data was imported into R and visualized using the ggplot2 package (**Main text Fig 2J**).

1. Paxinos, G. & Watson, C. *The Rat Brain in Stereotaxic Coordinates*. (Elsevier, 2005).
